# Atypical chlorotic plants as a tool for studying more severe Cd effect on photosystem I, non-photochemical quenching, K content, and stomatal conductance

**DOI:** 10.64898/2026.01.30.702819

**Authors:** Eugene A. Lysenko, Natalia A. Savvina, Alexander V. Kartashov, Galina V. Kochetova

**Affiliations:** Institute of Plant Physiology RAS, 35 Botanicheskaya St., Moscow, 127276, Russia; Faculty of Biology, M.V. Lomonosov Moscow State University, Moscow 119234, Russia

**Keywords:** barley, cadmium, chlorophyll fluorescence, chlorosis, non-photochemical quenching, P_700_ light absorption, potassium, photosystem I, stomatal conductance

## Abstract

Cadmium is a very toxic heavy metal. We studied Cd-treated barley plants with especial focus on rare atypical plants with signs of chlorosis. Cd treatment decreased the maximal photochemical activities of both photosystems while the activity of photosystem I decreased more than activity of photosystem II. In photosystem II, Cd treatment inhibited non-photochemical quenching that increased portion of unquenched “closed” complexes of photosystem II. The latter effect increased balance of limitations between the acceptor side of photosystem II (qC) and the donor side of photosystem I (Y(ND)) and raised the ratio qC/Y(ND). All these effects were enhanced in the atypical more damaged plants. Cd treatment reduced K content in the first leaves; in atypical plants, K content decreased even more. Cd treatment changed a pattern of stomatal conductance possibly by means of reducing K content in leaves. The untreated barley plants kept different stomatal conductance at adaxial and abaxial sides of leaves and fulfilled a complicated diurnal dynamics with large ups and downs of stomatal conductance. The typical Cd-treated plants were less flexible and demonstrated medium values. Stomatal conductance in the untreated plants were higher or lower than in the typical Cd-treated plants depending on a particular time; average daytime stomatal conductance was equal in both variants. At 10.00, stomatal conductance in the atypical Cd-treated plants was smaller than in the typical ones. Levels of 13 chloroplast mRNAs remained unchanged, while *psbD* decreased in both types of Cd-treated plants.

**Highlights:**

- Several Cd effects were enhanced in more damaged (atypical) chlorotic plants
- Cd treatment decreased activity of photosystem I and non-photochemical quenching
- Ratio of limitations between photosystems II and I [qC/Y(ND)] was rather constant
- Cd treatment reduced K content in the first leaves
- Cd treatment changed pattern of stomatal conductance

## 1. Introduction

Cadmium is a heavy metal with high toxicity. It is widespread all around the world. Many territories have high Cd content (Pan et al. 2010; Zou et al. 2021) of both natural and anthropogenic origin. Cadmium is rather mobile in natural conditions (Gaillardet et al. 2003, Jigyasu et al. 2020); plants absorb Cd by roots from soil. Most of the terrestrial plants restrict Cd movement from roots to shoots; they try to exclude Cd accumulation in aboveground part, therefore, they called excluders (Baker 1981).

Chloroplasts are important target for Cd action. Chloroplasts generate energy for both growth and protective mechanisms. Plants need to protect their chloroplasts from Cd accumulation. *In vivo*, small amount of Cd invades chloroplasts (Baryla et al. 2001; Pietrini et al. 2003; Lysenko et al. 2015). Inside chloroplasts, Cd is mainly located in thylakoids while its content in stroma is low (Lysenko et al. 2019). In thylakoids, Cd content is close to the content of Cu and comparable to the contents of Zn and Mn; therefore, Cu-dependent plastocyanin, Zn-dependent carbonic anhydrases and FtsH proteases, and Mn-dependent oxygen-evolving complex (OEC) of the photosystem II (PSII) can be considered as the target for the inhibition by Cd substitution of corresponding cations (Lysenko et al. 2019). Cadmium inhibition of OEC is generally recognized. *In vitro*, Cd inhibited OEC, while enormous Cd concentrations applied in these experiments (Bazzaz, Govindjee 1974; Sigfridson et al. 2004; Faller et al. 2005). In intact chloroplasts, such high Cd concentrations can be achieved neither *in vivo*, nor even *in vitro* (Lysenko et al. 2015; 2019). *In vivo*, the only experiment showed that Cd inhibited PSII while the photosystem I (PSI) remained rather uninfluenced (Baszynski et al. 1980); the same demonstrated *in vitro* (Bazzaz, Govindjee 1974). However, Baszynski et al (1980) demonstrated drastic decrease of phylloquinone (vitamin K_1_) that is the key component of PSI. Siedlecka and Baszynski (1993) demonstrated Cd inhibition of PSI *in vivo*. PSI demonstrated resistance to Cd action when the artificial electron acceptor methyl viologen was used (Baszynski et al. 1980; Siedlecka and Baszynski 1993); Cd inhibition of PSI *in vivo* revealed with use of the natural electron acceptor NADP only. This implies that Cd does not damage PSI itself but inhibits a component in close vicinity to PSI. Siedlecka and Baszynski (1993) pointed to ferredoxin.

In our experiments, Cd treatment inhibited PSI more than PSII; this was shown with the ratios of the photochemical activities in both dark-adapted (Pm/Fv) and light-adapted (Y(I)/X(II)) conditions (Lysenko et al. 2020). Cd treatment decreased non-photochemical quenching and increased acceptor-side limitation of PSII (Lysenko et al. 2020), while OEC functions at the donor-side of PSII. These results suggest that *in vivo* Cd inhibits the electron-transfer chain between PSII and PSI. We hypothesized Cd/Cu substitution in plastocyanin as the site of inhibition (Lysenko and Kusnetsov 2024). We tried to increase the observed effects; however, it was not so easy. The above mentioned results obtained in nine-day old barley plants at 80 μM Cd; this concentration is toxic but far from lethal (Klaus et al. 2013; Lysenko et al. 2015). Concentration 250 μM Cd is lethal in prolonged experiments (Klaus et al. 2013; Lysenko et al. 2015); at 250 μM Cd both barley and maize plants are severely depressed. However, barley leaves accumulated less Cd at 250 μM Cd than at 80 μM Cd; Cd content in chloroplasts also decreased while the decrease was insignificant (Lysenko et al. 2015). Therefore, neither photochemical effect enlarged at 250 μM Cd (Lysenko and Kusnetsov 2024). The nine-day old maize plants demonstrated different pattern of the changes. Solely, Pm/Fv was unchanged at 80 μM Cd and decreased at 250 μM Cd (Lysenko and Kusnetsov 2024). We have found another model to reveal increasing of Cd effect. Among typical Cd-treated plants, we analyzed rare (atypical) plants with more damaged phenotypes; more damaged plants demonstrated larger Cd effect on the processes around PSII and PSI.

We studied plants grown at 80 μM Cd. Accidentally, we observed three unusual maize plants with necrosis along veins in the first leaves. In these atypical plants, PSI (Pm) decreased more than PSII (Fv) comparing with the typical plants from the same pot. We continued the investigation with barley because barley plants accumulated more Cd in their chloroplasts (Lysenko et al. 2015) and demonstrated larger changes of PSI and PSII activities (Lysenko and Kusnetsov 2024). Barley plants demonstrated no necrosis along leaf veins; barley plants with tip, edge, or spot necrosis showed no more decrease of Pm/Fv. Using barley plants, we had to extend Cd treatment by one more week and study atypical plants with signs of chlorosis instead of necrosis. We have studied 16-day old barley plants grown at 80 μM Cd and compared typical and atypical plants from the same vessels. We measured maximal and actual activities of PSII and PSI using Dual-PAM-100. We have shown the enlargement of the previously found Cd effects on the processes around PSI and PSII (Lysenko et al. 2020) in more damaged plants. We have compared the results obtained with PAM and low temperature spectra of chlorophyll (Chl) fluorescence. We have studied the contents of Cd and essential metals, diurnal dynamics of stomatal conductance, and levels of chloroplast mRNAs. For the first time, we have shown a correlation between Cd effect, K content and stomatal conductance. The results are presented below.

## 2. Material and methods

### 2.1. Plant growth conditions

Barley (*Hordeum vulgare* L. cv. Luch) seedlings were grown in phytotron chambers at 21°C, 180–220 µmol photons m^−2^ s^−1^ and a photoperiod of 16 h light/8 h dark on modified Hoagland medium under continuous aeration (Lysenko et al. 2019). Preliminary, caryopses were kept for 2–3 days at 4 °C in the dark on a filter paper moistened with 0.25 mM CaCl_2_; imbibed caryopses were germinated in the growth conditions. Two-day old seedlings were transferred to vessels with Hoagland medium. The next day, CdSO_4_ was added to the hydroponic media to the final concentrations 80 µM. The Hoagland medium was renewed once a week. All the analyses were performed on the first leaves of 16-day old seedlings. All experiments repeated at least three times. For more details see (Lysenko et al. 2015; 2019; 2020). Maize (*Zea mays* L. cv. Luchistaya) seedlings were grown once under the same conditions but under 25-26℃.

### 2.2. Selection of atypical plants

On the vessels containing 80 µM Cd, green leaves with minimal damages were selected as typical (Lysenko et al. 2015; 2019); leaves with yellowish shade were chosen for further analysis. In all leaves, Fv and Pm were measured. Plants with yellowish shade and Fv/Pm ≥ 3.8 were selected as atypical. The threshold level Fv/Pm = 3.8 (Pm/Fv = 0.263) selected empirically. In the early experiments, the typical Cd-treated plants showed the ratio Fv/Pm < 3.6; later on, 3 out of 76 typical plants showed Fv/Pm = 3.9-4.1.

We were greatly limited with the number of atypical plants. The diurnal dynamic of stomatal conductance was not studied in the atypical plants. For the measurement of low temperature Chl fluorescence, we used atypical plants including those with Fv/Pm < 3.8 (up to 3.1); three curves were similar to the majority of atypical plants (Fv/Pm > 3.8); single curve (plants with Fv/Pm 3.65 and 3.46) was drastically different and therefore discriminated. In 1 out of 4 mRNA experiments, two leaves (Fv/Pm > 3.8) were supplemented with two leaves (Fv/Pm = 3.52).

### 2.3. PAM analysis

Chl *a* fluorescence and P_700_ light absorption were simultaneously registered with the Dual-PAM-100 (Walz, Germany). The Chl *a* fluorescence was excited at 460 nm (Int. #5, 12 μmol photons m^−2^ s^−1^; the measuring light). P_700_ is the reaction center (RC) Chl of PSI; the level of oxidized P_700_ was measured as the difference in light absorption at 830 and 875 nm (Int. #5). The red light 635 nm was used as the actinic light (AL) and for the saturation pulses (SPs).

The plants were adapted in the dark for 30 min, transferred rapidly to the Dual-PAM-100, avoiding bright light, and kept in the dark for another 4 min. In the dark, the measuring light only was used for the determination of minimal (Fo) Chl fluorescence; the first SP (4 mmol photons m^−2^ s^−1^, 500 ms) was applied for the measurement of maximal (Fm) Chl fluorescence; the pre-illumination with far-red light (720 nm, Int. #10, ∼250 μmol photons m^−2^ s^−1^, Pfündel et al. 2013) followed with the second SP were employed for the determination of minimal (Po) and maximal (Pm) P_700_ absorption. The measurement of basic parameters in the dark-adapted state was followed with the extra 30 s of darkness.

Next, the AL 168 μmol photons m^−2^ s^−1^ was induced for 7.5 min and induction curves (ICs) were recorded; SPs (4 mmol photons m^−2^ s^−1^, 500 ms) were induced every 40 s and followed by far-red light (720 nm, Int. #10, ∼250 μmol photons m^−2^ s^−1^, Pfündel et al. 2013) for 5 s.The maximum level of fluorescence in the light (Fm’) and maximal P_700_ change in the light (Pm’) were measured with SP; the stationary Chl fluorescence (Fs, synonym F) and P_700_ light absorption (P) were measured just before SP (Pfündel et al. 2013). After SP, the illumination was switched to the far-red light (720nm, Int. #10, ∼250 μmol photons m^−2^ s^−1^, Pfündel et al. 2013) for 5 s and the minimum level of fluorescence in the light (Fo’) was determined. The minimal level of P_700_ light absorption (Po) was measured after cessation of far-red light both in the dark and light regimes (Klughammer and Schreiber 1994; 2008).

The IC method is the correct and, probably, preferable way of plant adaptation to light conditions before an application of rapid light curve (RLC) method (Lysenko 2021). After the completing of IC, the RLC method was started in 6 s (software’s switching time). In RLC, each step of AL lasted for 30 s; the set of AL intensities is shown in the figures. The other parameters were the same as in IC. The RLC method aspires to but does not reach a stationary level (White and Critchley 1999; Kalaji et al. 2014); RLC data should be considered as quasi-stationary.

The parameters of P_700_ light absorption were calculated as follows: Y(I) = (Pm’ – P)/(Pm – Po); Y(ND) = (P – Po)/(Pm – Po); Y(NA) = (Pm – Pm’)/(Pm – Po); Y(I) + Y(ND) + Y(NA) = 1 (Klughammer and Schreiber 1994; 2008).

The parameters of Chl fluorescence were calculated using the following equations: Fv = Fm – Fo; Fv’ = Fm’ – Fo’; Φ_PSII_ = (Fm’ – Fs)/Fm’ (Genty et al. 1989; Kalaji et al. 2014); ΔF = Fm’ – Fs (Lichtenthaler et al. 2005); ΔC = Fs – Fo’, ΔF + ΔC = Fv’ (Lysenko 2021); qP = (Fm’ – Fs)/Fv’, qN = (Fv – Fv’)/Fv (Schreiber et al. 1986; van Kooten and Snel 1990); X(II) = (Fm’ – Fs)/Fv, qC = (Fs – Fo’)/Fv, qN + X(II) + qC = 1 (Lysenko et al. 2020).`#`

The ratios qC/Y(ND) calculated according to (Lysenko and Kusnetsov 2024). In atypical plants, 27 RLC were collected; among them, 3 datapoints discriminated at 72 μmol photons m^−2^ s^−1^and single datapoint discriminated at 128 μmol photons m^−2^ s^−1^ (not shown). In atypical plants, many values Y(I) and X(II) were close to zero in RLC at AL > 432 μmol photons m^−2^ s^−1^; hence, too many datapoints of the ratio Y(I)/X(II) must be discriminated. Therefore, the dynamics of Y(I)/X(II) ratio at 662–1954 μmol photons m^−2^ s^−1^ is not shown in atypical plants.

### 2.4. Low-temperature (77K) fluorescence spectroscopy

After Fv and Pm determination, plants were adapted to the dark at room temperature for 1 h. We wrapped the plants with wet filter paper and kept on ice in darkness (for a transfer from IPP RAS to Lomonosov MSU). For each sample, 2-cm leaf segments were collected from three typical plants (Cd 0 µM or Cd 80 µM) or 3-cm leaf segments from two atypical plants (Cd 80 µM); rarely, a sample was prepared from 6-cm leaf segment of a single atypical plant (Cd 80 µM). The whole procedure performed according to Weis (1985) and Kochetova et al (2018). Tissues were grinded with pestle and mortar in the medium (25 mM Tris-HCl, pH 8.0; 350 mM NaCl, 5 mM MgCl_2_) on ice for about 1.5 min. The homogenate was filtered through a nylon cloth with the mesh size of ∼50 μm; glycerol was added to the final concentration of 65% (v/v). Each aliquot of 75 µl was carefully frozen in liquid nitrogen. The transparent 1.5 mm wide samples were measured. The fluorescence emission spectra were measured in liquid nitrogen (77K) with a FluoroMax-4 spectrofluorometer (HORIBA-Scientific, France, Lomonosov MSU Shared User Facility «FluoroMax-4 spectrofluorometer for fluorescence spectroscopy»). The fluorescence was excited at 440 nm; the emission was recorded in between 650 and 800 nm. The width was 2 nm for both entrance and exit slits. The background (blank vial) was subtracted; the spectra were normalized to the maximum at 683 nm, smoothed by Savitzky-Golay method and averaged using OriginPro 2015 (OriginLab Corporation, USA).

### 2.5. Measurement of metals

The dry leaves were digested with nitric and perchloric acids as described in (Lysenko et al.2020). The metal contents were determined with a flame atomic absorption spectrophotometer (AA-7000, Shimadzu, Japan) with a hollow-cathode lamps (Hamamatsu, Japan). Prior to the analysis, samples were appropriately diluted with 0.1M HNO_3_. For the measurement of Ca and K, the diluting solutions included LaCl₃ (10 g L⁻¹) or LiCl (1 g L⁻¹) respectively to mitigate ionization interference.

### 2.6. Measurement of stomatal conductance

The stomatal conductance was measured with the porometer/fluorometer Li-600 (Li-Cor Biosciences, USA) according to the manufacturer’s instruction. The conductance was defined as g_sw_ = H_2_O mol m^−2^ s^−1^. A single leaf was measured in three points from one side (adaxial or abaxial) that were averaged in a single datapoint for further calculations.

### 2.7. RNA extraction and RT-PCR

Mortar, pestle and leaves were chilled with liquid nitrogen; leaves were grinded with the RNA extraction buffer (20 mM Tris-HCl, pH 7.0, 4 M guanidine thiocyanate, 20 mM EDTA, 0.7% lauroyl sarcosin, 100 mM β-mercaptoethanol; 1 : 2 (m/v)) according to Lysenko et al. (2013). An RNA sample was extracted from 4±1 first leaves. Residual DNA removed by DNase I (Thermo Fisher Scientific, USA) treatment. The first cDNA strand was synthesized with the reverse transcriptase Maxima H Minus (Thermo Fisher Scientific, USA) using random decamer primers (dN_10_, Lytech, Russia). The cDNA synthesis performed with 2.25 μg of total RNA, 100 U of the reverse transcriptase, and ∼100 pmol of primers in 25 μl; the samples pre-incubated at 25°C for 7 min for primer annealing and synthesis performed at 60°С for 45 min. A cDNA from 45 ng of total RNA was amplified with Taq DNA polymerase in 20 μl according to the manufacturer’s recommendations (ABclonal, China). For each gene, a cDNA specimen was amplified at least twice with different number of cycles. Sequences of primers, annealing temperatures and numbers of amplification cycles are given in Suppl Table S1. PCR products were analyzed by electrophoresis in a 1.5% agarose gel with ethidium bromide and results were registered using iBright CL1500 imaging system (Thermo Fisher Scientific, USA). Bands of psbD PCR-products were deciphered with software in iBright CL1500.

### 2.8. Statistics

Each experiment performed in three independent experiments at least. As a whole, the thirteen independent biological experiments performed and ∼2,500 Cd-treated plants analyzed (Suppl Table S2).

The data processed using the Excel (Microsoft) software. Means are mostly represented by arithmetic means. Once geometric means were calculated, it was specified particularly. The significance of differences between mean values verified with *t*-test.

## 3. Results

In a routine experiment, we studied nine-day old maize plants grown at 80 μM Cd. Accidentally, we observed three unusual maize plants with necrosis along veins in the first leaves. In typical Cd-treated maize plants, Pm decreased more than Fv. In three atypical plants, Pm and Pm/Fv decreased more comparing with the typical Cd-treated plants from the same vessel (Fig. 1). This suggests plants with a heavily damaged phenotype as a tool for the investigation of Cd effect on the photosystems *in vivo*. We have chosen barley to continue the experiment because barley chloroplasts accumulated more Cd comparing with maize chloroplasts (Lysenko et al. 2015).

**Figure 1.**
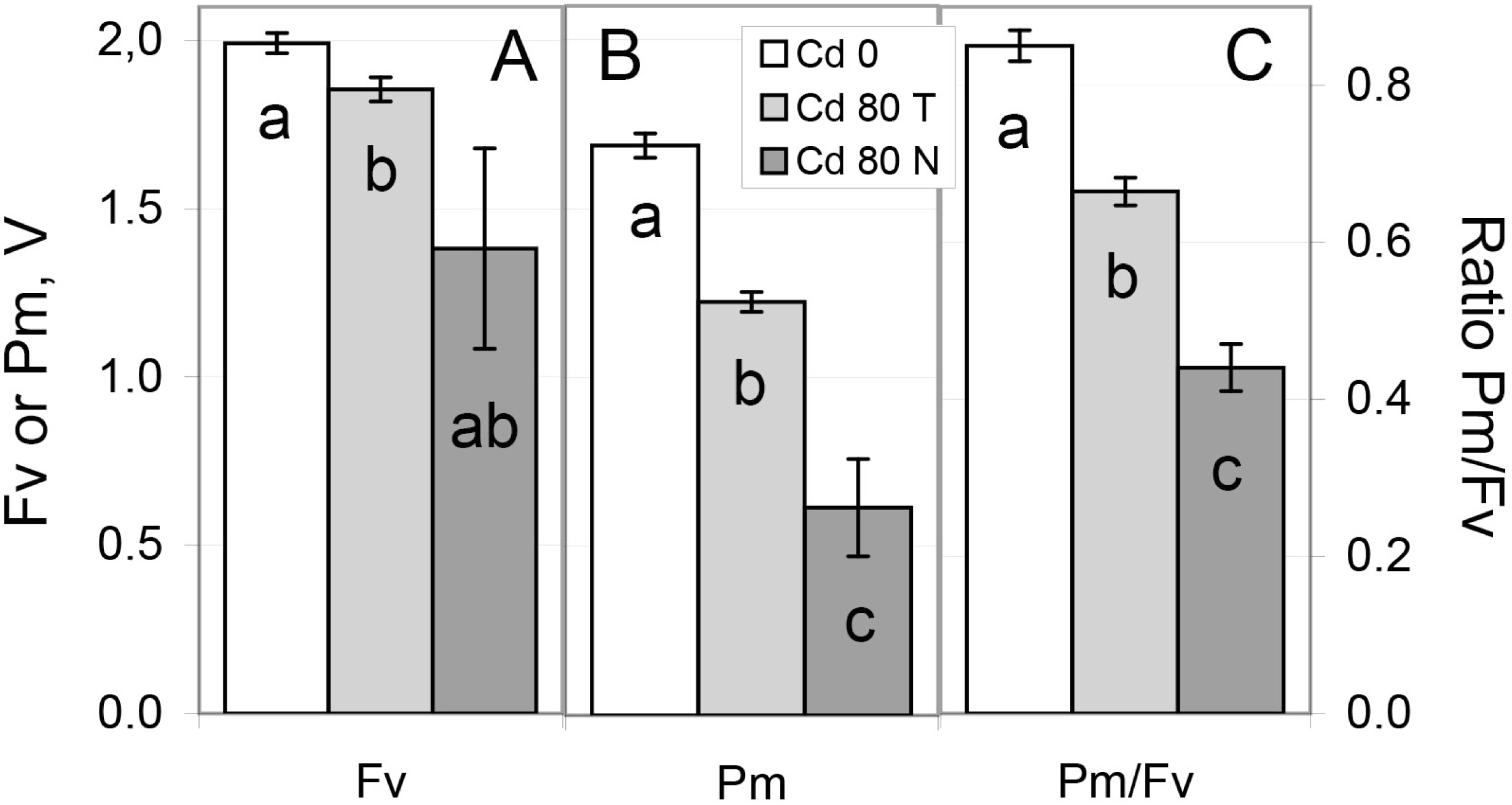
The maximal photochemical values in the dark-adapted 9-day old maize plants. A – Fv, B – Pm, C – the ratio Pm/Fv. White bars – untreated plants with no Cd addition (Cd 0); light grey bars – typical plants at 80 μM Cd (Cd 80 T); dark grey bars – atypical plants with necrosis along leaf veins at 80 μM Cd (Cd 80 N). The typical and atypical plants were grown on a same vessel at 80 μM Cd. Means ± standard errors (SE) are shown. a-c – the differences are significant at p ≤ 0.05.

We had to extend Cd treatment by one more week to obtain small but reproducible amount of atypical barley plants. At 80 μM Cd, 16-day old barley plants showed no necrosis along leaf veins; plants with other types of necrosis and apparent local chlorosis demonstrated no especial decrease of Pm comparing with Fv. The further Pm decrease was found in barley plants with extensive chlorosis. Typically, nor 9-day old (Lysenko et al. 2015; 2020), neither 16-day old (Fig. 2) barley plants demonstrated signs of chlorosis at 80 μM Cd. Among the typical plants, we found atypical plants with an extensive yellowish shade on the first leaf (Fig. 2B); such atypical plants comprised 2-3% of all the plants at the vessel with 80 μM Cd (Suppl. Table S2). This study focuses on the comparison of the typical and atypical (chlorotic) plants grown at same vessels at 80 μM Cd. The typical Cd-treated plants (Cd80 Typical) are the first control for the atypical Cd-treated plants (Cd80 Atypical). The typical plants grown with no Cd addition (Cd0 or untreated) are the second control for a general Cd effect. At the 16^th^ day, the untreated plants were much larger than Cd-treated ones (Fig. 2A). This large difference in the growth and development itself can induce differences between inner processes in Cd-treated and -untreated plants. Therefore, we compared the untreated (Cd0) and Cd-treated (Cd80 Typical and Atypical) variants with a caution. In the most experiments, atypical leaves were moderately damaged (Fig. 2A) while severally damaged leaves obtained in a sole experiment (Suppl. Fig. S1); a completely yellow first leaf was obtained once (not shown).

**Figure 2.**
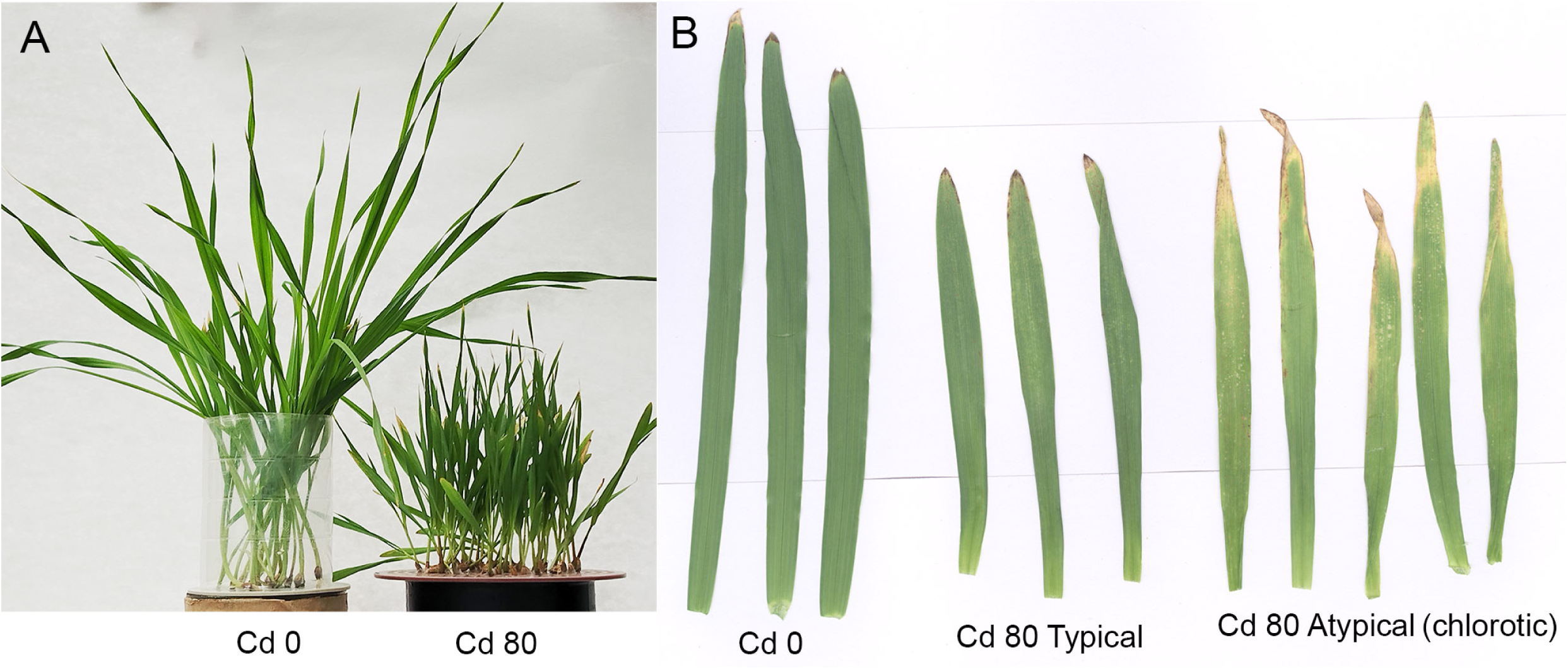
The phenotypes of 16-day old barley plants. A – 16-day old barley plants; B – the first leaves of 16-day old barley plants. Cd 0 – untreated plants with no Cd addition; Cd 80 – plants grown at 80 μM Cd. Both typical and atypical plants were grown on same vessels at 80 μM Cd. Part B shows common phenotype of atypical leaves; the most damaged atypical leaves are shown in Suppl. Fig. S1.

### 3.1. The maximum activities of PSII and PSI in the dark-adapted state

The maximum values of Chl fluorescence and P_700_ light absorption are shown in Fig. 3. In the dark-adapted state, the fluorescence of antennae (Fo) and RCs (Fv) of PSII and the redox changes of P_700_ (Pm) decreased in the typical Cd-treated plants and much more decreased in the atypical ones (Fig. 3A–C). In the typical Cd-treated plants, these values felt nearly proportionally and their ratios Fv/Fm and Pm/Fv did not change (Fig. 3D–E). In the atypical Cd-treated plants, Pm reduced much more than Fv while Fv decreased slightly more then Fo; therefore, the ratio Pm/Fv decreased more than twofold while the ratio Fv/Fm decreased slightly (Fig. 3D–E). Some atypical plants demonstrated very low values of Pm and Fv; accordingly, the smallest and largest values differed by an order of magnitude (Fig. 3F). Therefore, we calculated both arithmetic and geometric means for all these values; they were indistinguishable (Fig. 3A–E). The individual Pm/Fv ratios showed a cloud-like distribution in the typical Cd-treated plants, a weak tendency in the untreated plants and a strong mutual decrease of Pm and Fv in the atypical plants (Fig. 3F); plants with light signs of chlorosis positioned between the typical Cd-treated plants and the atypical plants with apparent signs of chlorosis (Fig. 3F).

**Figure 3.**
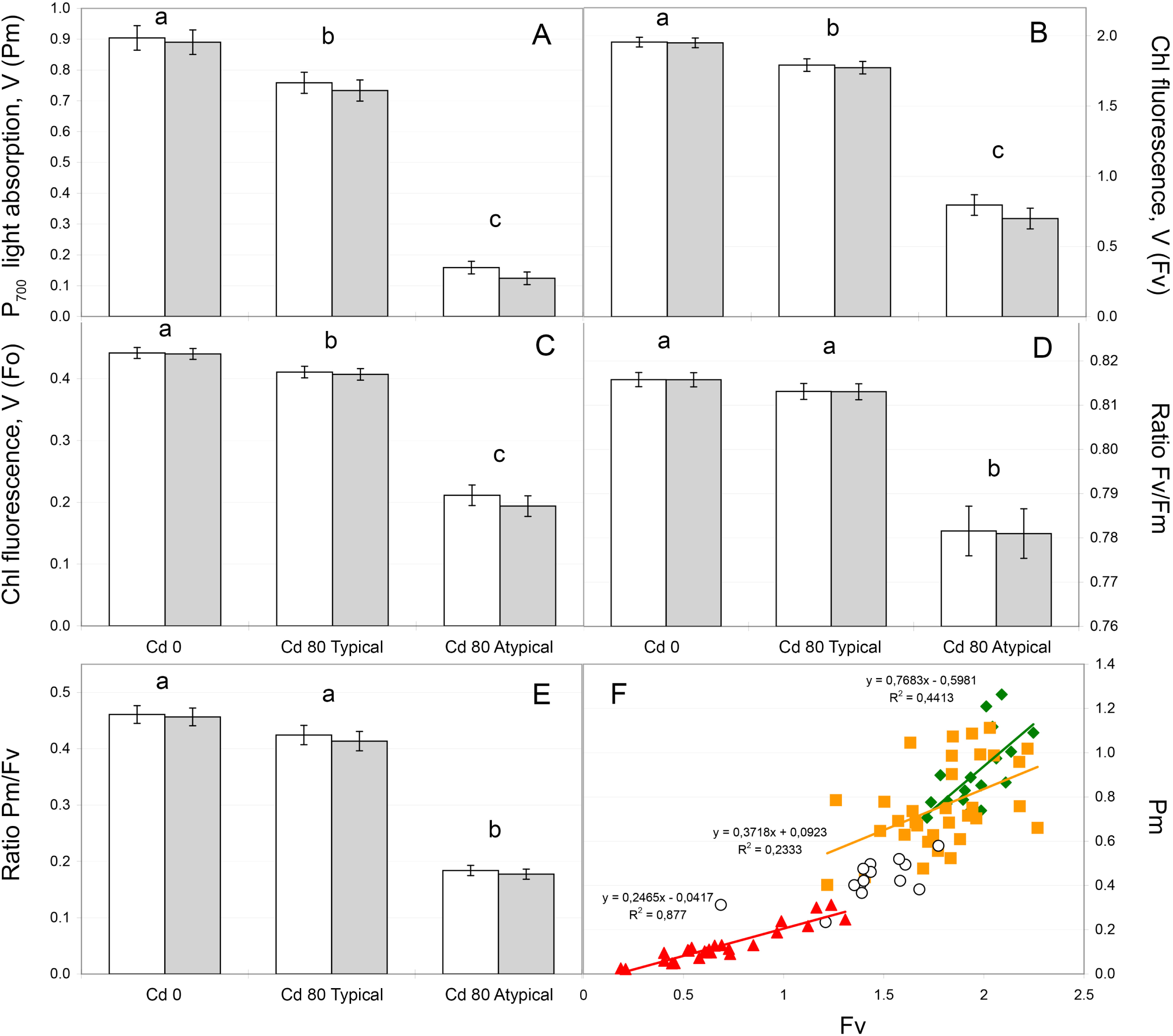
Basic photochemical values in the dark-adapted 16-day old barley plants. A – Pm, B – Fv, C – Fo, D – the ratio Fv/Fm, E, F – the ratio Pm/Fv. Cd 0 – untreated plants with no Cd addition; Cd 80 – plants grown at 80 μM Cd, both typical and atypical with signs of chlorosis. A–E: white bars – arithmetic means, grey bars – geometric means. Means ± standard errors (SE) are shown. a-c – the differences are significant at p ≤ 0.05. F: green diamonds – untreated plants (Cd 0), orange squares – Cd-treated typical plants (Cd 80 Typical), white circles – Cd-treated atypical plants with light chlorosis (mostly discarded), red triangles – Cd-treated atypical plants with distinct chlorosis (Cd 80 Atypical).

### 3.2. Activity of PSII under illumination

Under light conditions (AL), the photochemical activity of PSII increased in the typical Cd-treated plants according to any coefficient: ΔF, ΔF/Fv (X(II)), ΔF/Fv’ (qP), or ΔF/Fm’ (Φ_PSII_) (Fig. 4A-B, Suppl. Fig. S2); solely, qP in RLC was nearly equal in the untreated and typical Cd-treated plants (Suppl. Fig. S2H). In the atypical plants, the maximum activity of RC PSII (Fv) drastically decreased (Fig. 3B); therefore, under AL conditions, ΔF in the atypical plants was smaller than in the untreated plants (Suppl. Fig. S2A–B). The result depended on a value to which ΔF was normalized. If the effective activity in light (ΔF) normalized to the maximum activity in darkness (X(II) = ΔF/Fv), then PSII activity in the atypical plants remained at the level of typical Cd-treated plants in IC (Fig. 4A) and decreased back to the level of untreated plants in RLC (Fig. 4B). If ΔF normalized to values also measured under AL conditions (qP = ΔF/Fv’, Φ_PSII_ = ΔF/Fm’), then PSII activity in the atypical plants felt below the level of untreated plants (Suppl. Fig. S2E–H). According to qP, this fall is large; however, it caused by an increase of closed PSII (ΔC = Fs – Fo’) because qP shows the balance between open (ΔF) and closed (ΔC) RC PSII and can be presented as qP = ΔF/(ΔF + ΔC).

**Figure 4.**
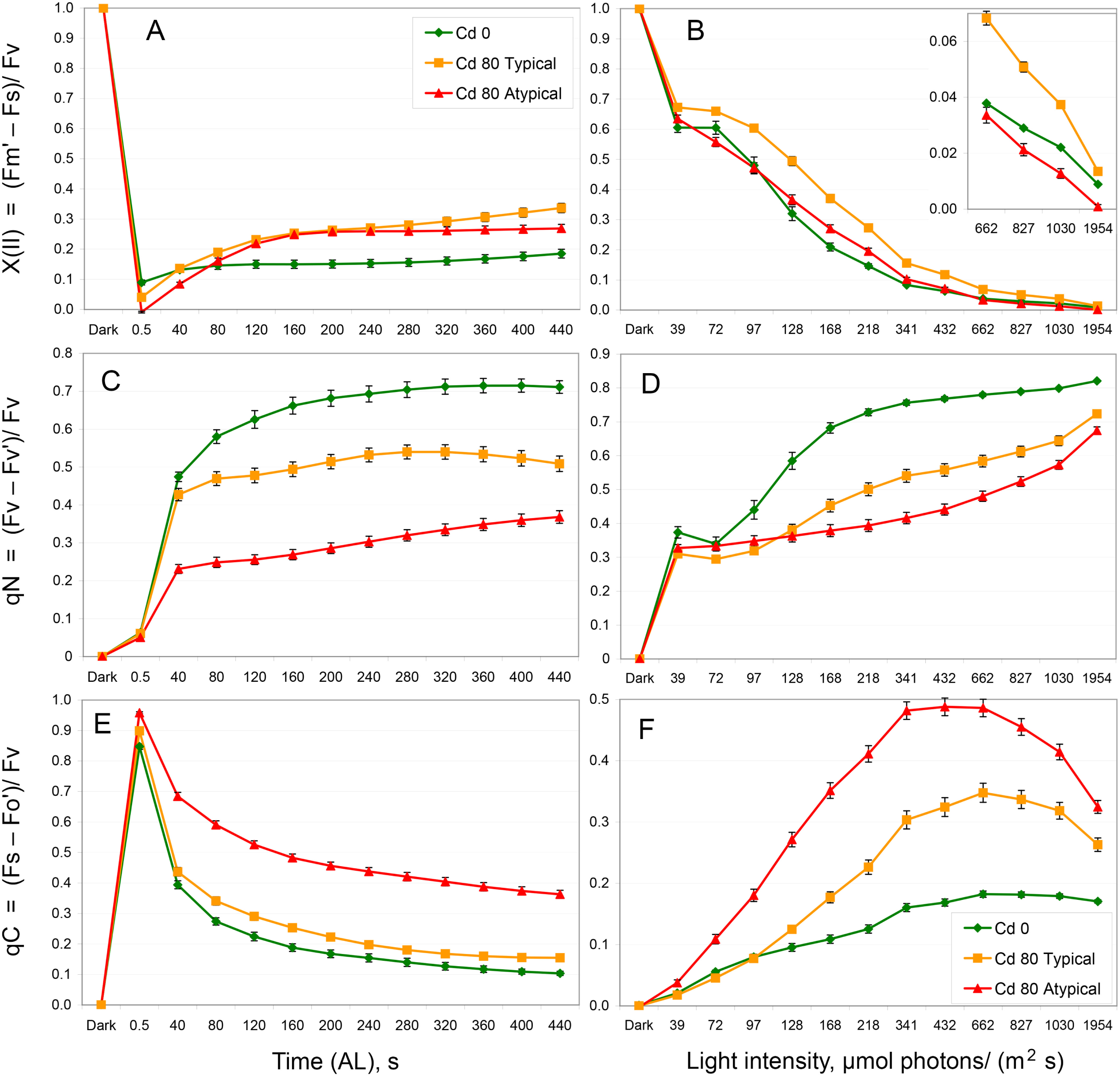
Dynamics of the photochemical and non-photochemical parameters of PSII under illumination in 16-day old barley plants. A, B – the photochemical coefficient X(II) (dynamics of ΔF and ΔF-based coefficients X(II), Φ_PSII_, and qP are given in Suppl. Fig. S2); C, D – coefficient of non-photochemical quenching qN; E, F – coefficient qC showing portion of closed RC PSII (dynamics of ΔC and related coefficients qC and 1–qP are given in Suppl. Fig. S3). A, C, E – IC; B, D, F – RLC. Green diamonds – untreated plants (Cd 0), orange squares – Cd-treated typical plants (Cd 80 Typical), red triangles – Cd-treated atypical plants (Cd 80 Atypical). Means ± SE are shown. The inset shows the corresponding small values with the higher resolution.

Cd-treatment decreased non-photochemical quenching by PSII in the typical plants; in more damaged atypical plants, the decrease was enhanced (Fig. 4C–D). As a result, the portion of closed RC PSII (qC = ΔC/Fv) increased in Cd-treated plants. In the typical Cd-treated plants, qC increased slightly in IC (Fig. 4E) and substantially in RLC (Fig. 4F). In more damaged atypical plants, qC was further increased in both IC and RLC comparing with the typical Cd-treated plants (Fig. 4E–F). The other coefficient 1–qP changed the shape of ΔC dynamics in RLC; in the typical Cd-treated plants, the coefficient 1–qP hid qC increase in RLC and turned slight qC increase to small qC decrease in IC (Suppl. Fig. S3). This is because qP solely shows the balance between opened and closed RC PSII and excludes non-photochemical quenching from consideration.

### 3.3. Activity of PSI under illumination

Under light conditions (AL), the photochemical activity of PSI also increased in the typical Cd-treated plants (Fig. 5A–B). In the atypical plants, Y(I) decreased back to the level of untreated plants or below it (Fig. 5A–B). This was caused by the inverse changes of limitation at the donor-side of PSI (Fig. 5C–D). Limitation at the acceptor-side of PSI remained small (Fig. 5E–F). Cd treatment increased Y(NA) slightly; in the typical plants, Y(NA) increased significantly in the most points of IC and RLC; in the atypical plants, the increase was similar and significant in the most points of IC since 200 s.

**Figure 5.**
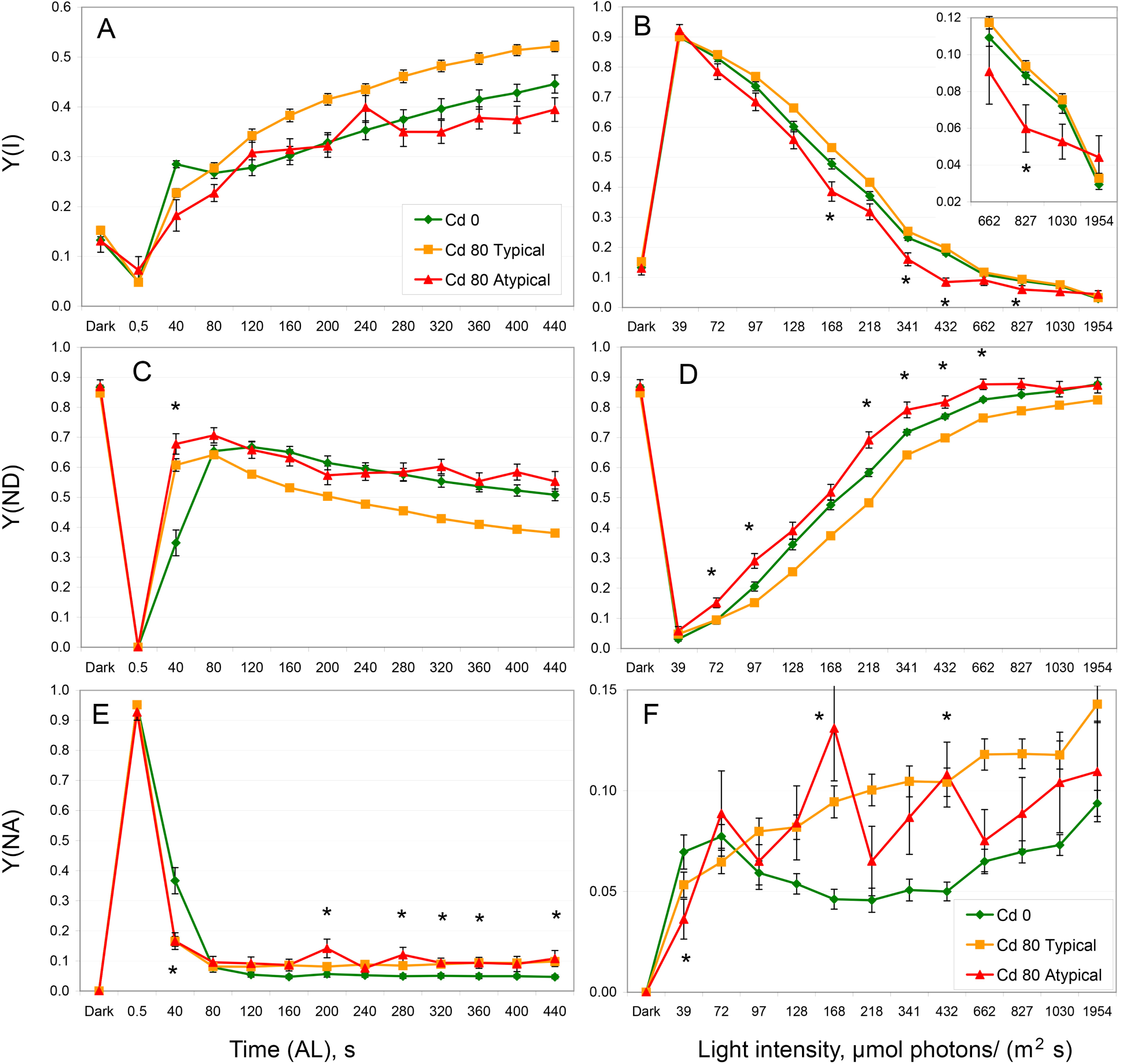
Dynamics of the photochemical activity and limitations of PSI under illumination in 16-day old barley plants. A, B – the photochemical coefficient Y(I); C, D – limitation of PSI at the donor side, Y(ND); E, F – limitation of PSI at the acceptor side, Y(NA); A, C, E – IC; B, D, F – RLC. Green diamonds – untreated plants (Cd 0), orange squares – Cd-treated typical plants (Cd 80 Typical), red triangles – Cd-treated atypical plants (Cd 80 Atypical). * marks significant differences between Cd 0 and Cd 80 Atypical, p ≤ 0.05. Means ± SE are shown. The inset shows the corresponding small values with the higher resolution.

### 3.4. The ratio of PSI/PSII activities and limitations qC/Y(ND)

Cd treatment decreased the ratio Y(I)/X(II) (Fig. 6A–B). In more damaged atypical plants, the ratio Y(I)/X(II) further decreased in the most of stationary points in IC and at 432 μmol photons m^−2^ s^−1^ in RLC (Fig. 6 A–B); at the higher AL intensities, data are not shown as questionable (see chapter 2.3).

**Figure 6.**
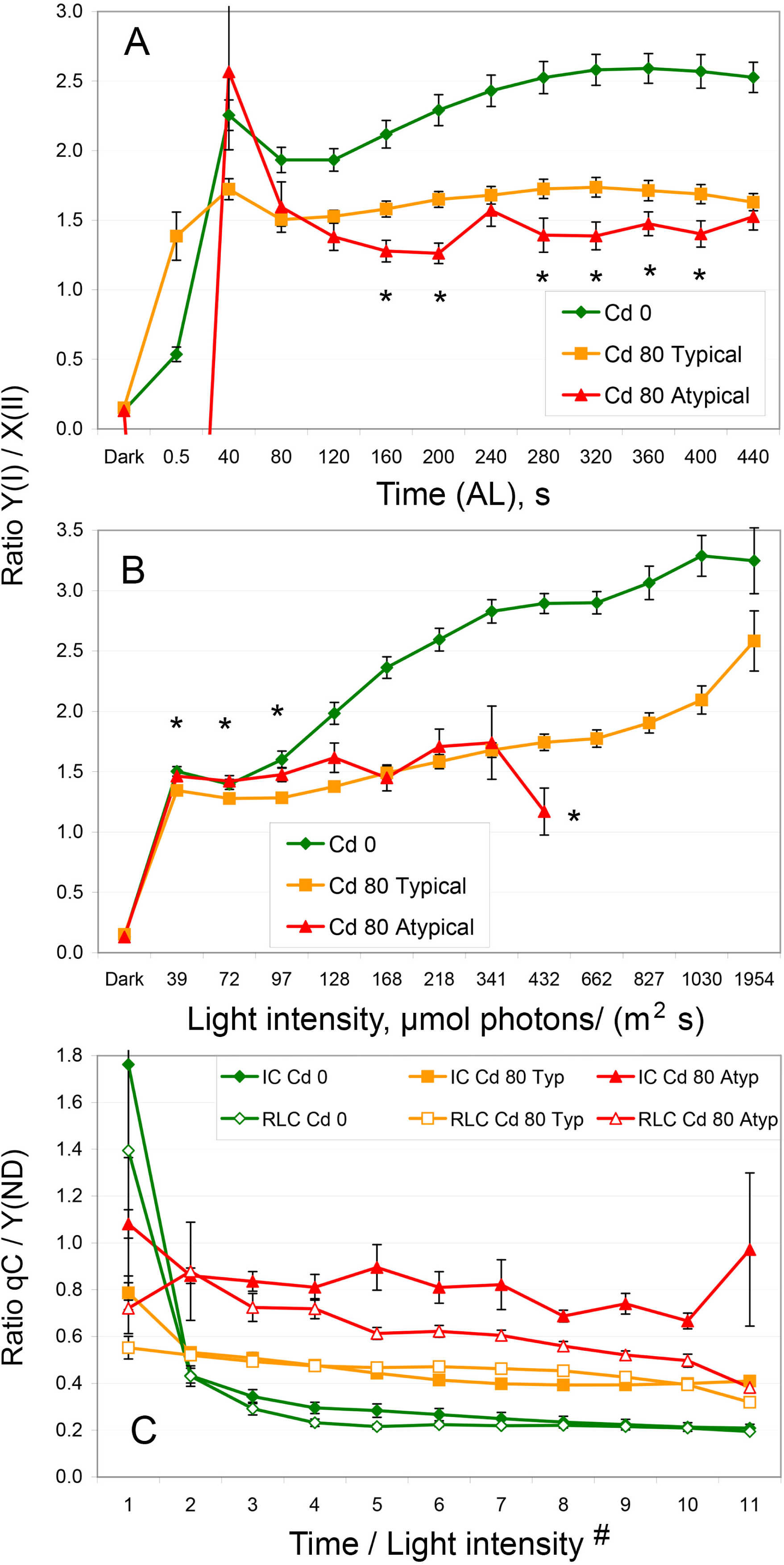
Balance of the photochemical activities (A, B) and limitations (C) between PSII and PSI in 16-day old barley plants. A, B – the ratio of photochemical activities of PSI and PSII, Y(I)/X(II). A – IC; B – RLC. Green diamonds – untreated plants (Cd 0), orange squares – Cd-treated typical plants (Cd 80 Typical), red triangles – Cd-treated atypical plants (Cd 80 Atypical). * marks significant differences between Cd-treated typical (Cd 80 Typical) and atypical (Cd 80 Atypical) plants, p ≤ 0.05. C – the ratio of limitations at the acceptor side of PSII and donor side of PSI, qC/Y(ND). Green diamonds – untreated plants (Cd 0), orange squares – Cd-treated typical plants (Cd 80 Typ), red triangles – Cd-treated atypical plants (Cd 80 Atyp); filled symbols show IC, open symbols show RLC. # – the abscissa numbering indicates both time and AL intensities. For IC, 1–11 correspond to the range 40 – 440 s (see A); for RLC, 1–11 correspond to the range 72 – 1954 μmol photons m^−2^ s^−1^ (see B). Means ± SE are shown.

Limitations at the acceptor side of PSII (qC) and donor-side of PSI (Y(ND)) changed with large amplitudes while their ratio remained rather constant (Fig. 6C). In untreated plants, the ratio qC/Y(ND) remained at the level ∼0.2 both in IC and RLC. In the typical Cd-treated plants, qC/Y(ND) increased to the level ∼0.4 both in IC and RLC. In more damaged atypical plants, qC/Y(ND) further rose to the level ∼0.8 in IC while gradually changed from 0.8 to 0.4 along with the growth of AL intensity in RLC (Fig. 6C).

### 3.5. Low temperature (77K) fluorescence spectra of Chl

We compared the results of PAM analysis with another experimental approach. The spectra of Chl fluorescence in liquid nitrogen and maximum values of Chl fluorescence and P_700_ light absorption were measured in the same plants (Fig. 7). In this experiment, Cd treatment decreased Pm more than Fv in the typical plants; in the atypical plants, Pm felt even more comparing with Fv (Fig. 7C–E). The spectra of low temperature Chl fluorescence were normalized to the peak of PSII at 683 nm (Fig. 7A). In contrast to the PAM data, PSI peak at 742 nm increased in the typical Cd-treated plants; in more damaged atypical plants, PSI peak at 742 nm decreased back to the level of untreated plants and slightly below it. The spans of significant differences mainly overlapped; however, the effect in typical Cd-treated plants was blue-shifted while the effect in atypical Cd-treated plants was red-shifted (Fig. 7B).

**Figure 7.**
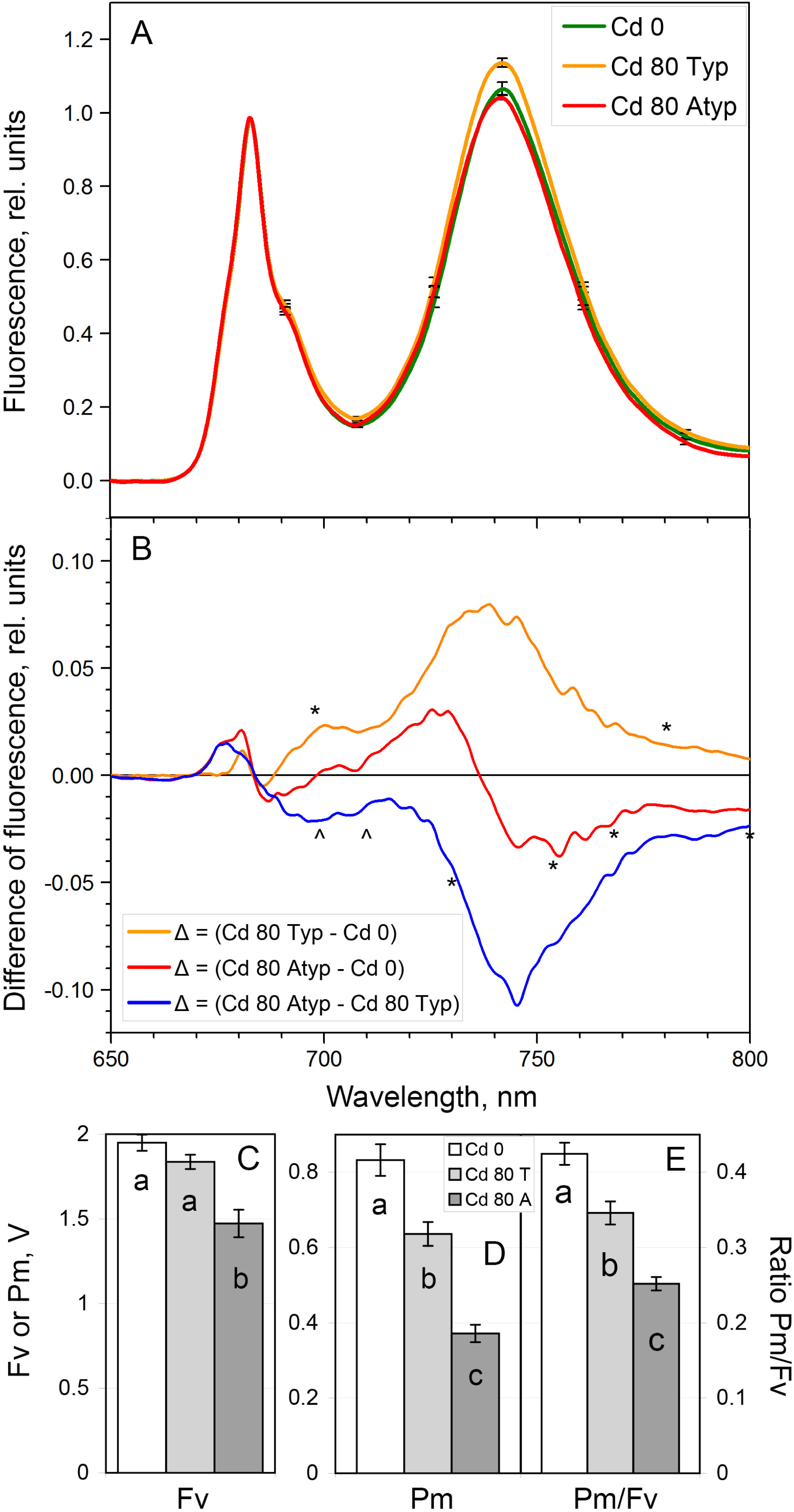
Low temperature (77K) Chl fluorescence in 16-day old barley plants. A – low temperature (77K) Chl fluorescence spectra normalized to the PSII maximum at 683 nm. Lines: green – untreated plants (Cd 0), orange – Cd-treated typical plants (Cd 80 Typ), red – Cd-treated atypical plants (Cd 80 Atyp). B – differences (Δ) between spectra from A: orange – [Cd 80 Typ] minus [Cd 0], red – [Cd 80 Atyp] minus [Cd 0], blue – [Cd 80 Atyp] minus [Cd 80 Typ]. Significance of the differences marked by symbols * and ^. Spans between two symbols * and between two symbols ^ are significant, p ≤ 0.05. C – Fv, D – Pm, E – Pm/Fv. White bars – untreated plants (Cd 0), light grey bars – Cd-treated typical plants (Cd 80 T), dark grey bars – Cd-treated atypical plants (Cd 80 A). Fv and Pm measured in the same leaves prior to the measurement of Chl fluorescence at 77K (A). Means ± SE are shown. a-c – the differences are significant at p ≤ 0.05.

### 3.6. Contents of Cd and essential metals in leaves

Were the signs of chlorosis caused by a higher Cd accumulation in leaves and/or chloroplasts? The amount of the atypical leaves was insufficient for chloroplast isolation; therefore, we solely measured Cd contents in leaves. The untreated plants demonstrated no Cd accumulation while Cd-treated plants showed large Cd content in the first leaves. The atypical plants demonstrated insignificant increase of Cd content comparing with the typical Cd-treated plants (Fig. 8A).

**Figure 8.**
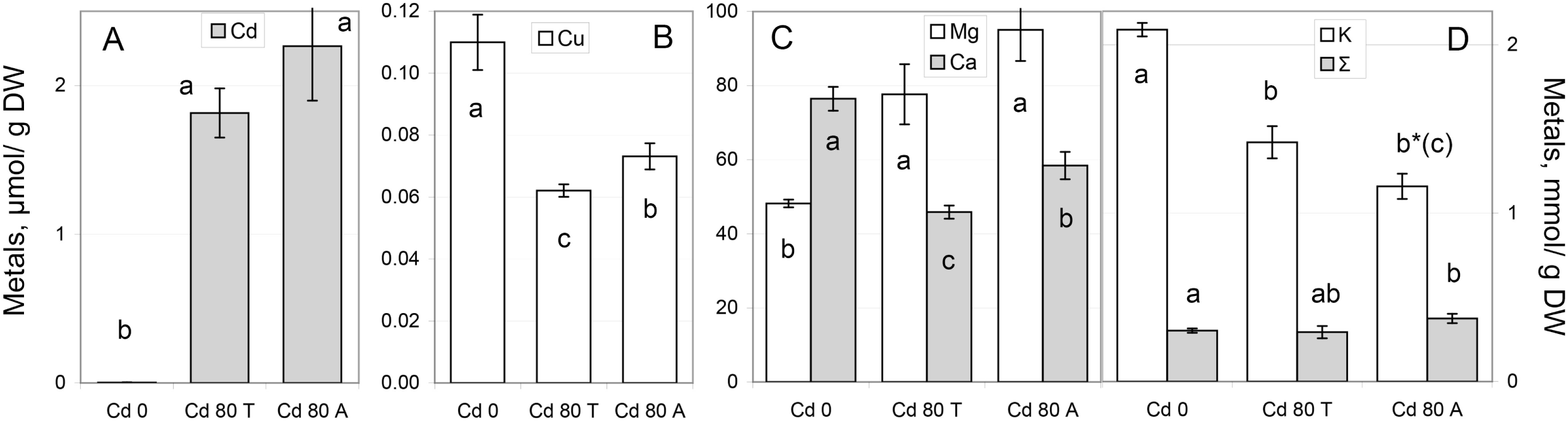
Metal contents in the first leaves of 16-day old barley plants. A – Cd; B – Cu; C – Mg (white) and Ca (grey); D – K (white) and Σ (grey). Σ – sum of all essential metals excepting K and some ultra-micro metals; Σ = [Na+Mg+Ca+Fe+Mn+Zn+Cu]. Cd 0 – untreated plants, Cd 80 T – Cd-treated typical plants, Cd 80 A – Cd-treated atypical plants. Means ± SE are shown. a-c – the differences are significant at p ≤ 0.05. * – difference between K contents in the typical and atypical Cd-treated leaves is significant at p < 0.055 per g DW and at p < 0.05 per a single leaf (Suppl. Table S3).

Essential metals showed remarkable changes. Potassium is the major metal in the barley first leaves (Fig. 8D). The content of K decreased in the typical Cd-treated plants and, probably, even more decreased in the atypical plants. The latter difference is significant per one leaf (Suppl. Table S3) and close to the significance per g DW (p < 0.055). Except K, the contents of all the other metals per g DW increased in the atypical plants comparing with the typical once whether significant or not (Suppl. Table S3). Two major bivalent cations demonstrated reciprocal changes: Cd treatment increased Mg while decreased Ca; in more damaged atypical plants, the contents of both metals increased comparing with the typical Cd-treated plants that was significant for Ca (Fig. 8C). The content of Cu changed similar to Ca: decreased in the typical Cd-treated plants and partially restored content in the atypical plants (Fig. 8B). The contents of Na, Fe, Mn, and Zn changed insignificantly (Suppl. Table S3).

### 3.7 Stomatal conductance

The large alteration of K content points to a possible change in stomatal conductance. We measured stomatal conductance at 10 a.m.; further, these plants were measured with PAM, transferred from IPP to MSU, and low temperature Chl fluorescence was determined. In morning, stomatal conductance (g_sw_) in the atypical first leaves was smaller than in the typical Cd-treated leaves; astonishingly, the stomatal conductance in the untreated plants also was smaller than in the typical Cd-treated plants (Fig. 9A). We tested the stomatal conductance throughout a day for both types of typical plants; all the atypical plants were measured at 10 a.m. and we had no extra plants to measure a diurnal dynamic.

**Figure 9.**
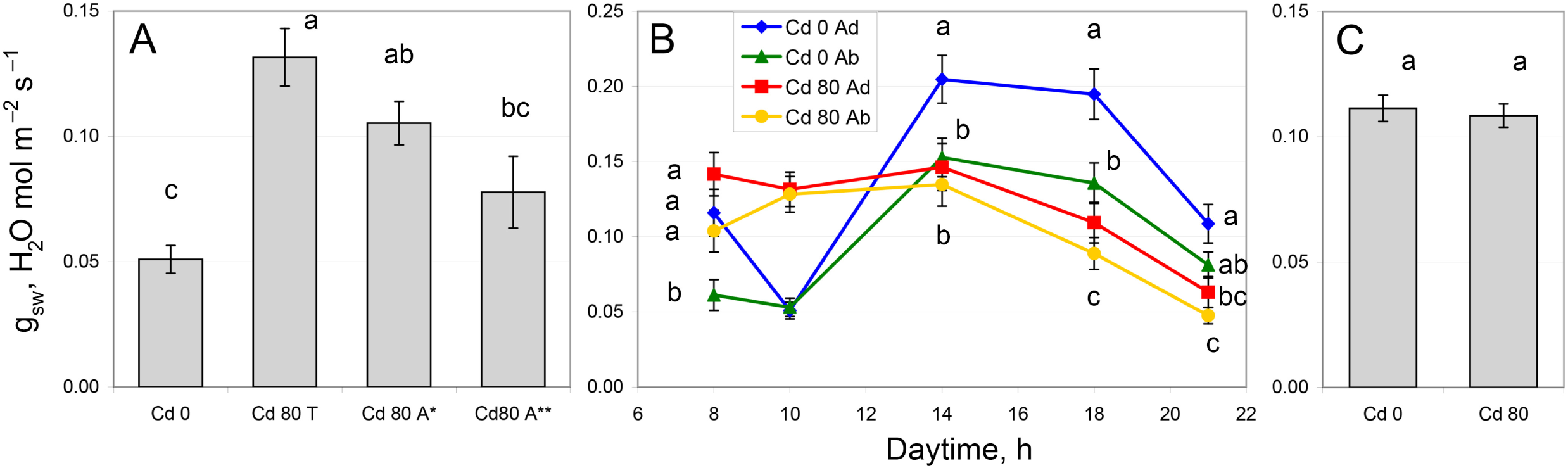
Stomatal conductance (g_sw_) in the first leaves of 16-day old barley plants. A – stomatal conductance at adaxial side of leaves measured at 10.00. Cd 0 – untreated plants, Cd 80 T – Cd-treated typical plants, Cd 80 A* – Cd-treated plants primary selected by phenotype (yellowish shade of the first leaf), Cd 80 A** – Cd-treated plants selected by phenotype and further demonstrated Fv/Pm ≥ 3.8 (Pm/Fv ≤ 0.263). B – diurnal dynamics of stomatal conductance; a day time from 7.00 to 23.00. Blue diamonds – adaxial (upper) side of untreated plants (Cd 0), green triangles – abaxial (lower) side of untreated plants (Cd 0), red square – adaxial (upper) side of Cd-treated typical plants (Cd 80), yellow circles – abaxial (lower) side of Cd-treated typical plants (Cd 80). C – average daytime stomatal conductance. For each variant, data for both sides of leaves measured at the five timeslots (see B) were averaged. Cd 0 – untreated plants, Cd 80 T – Cd-treated typical plants. Means ± SE are shown. a - c – the differences are significant at p ≤ 0.05.

The pattern of diurnal dynamic appeared different in the untreated and Cd-treated plants. The untreated barley plants changed the stomatal conductance extensively. They kept g_sw_ higher at the adaxial side of leaves than at abaxial side; they reduced g_sw_ at 10.00 and then greatly increased it at 14.00 (Fig. 9B). In Cd-treated plants, g_sw_ values were indistinguishable at both sides of leaves and from 10.00 till 14.00. A general feature was gradual decline of g_sw_ from 14.00 to 21.00 in both variants (Fig. 9B). The averaged daytime stomatal conductance was equal in both untreated and Cd-treated plants (Fig. 9C).

### 3.8. Chloroplast mRNA level

Finally, we tested chloroplast mRNAs. Previously, we demonstrated the absence of Cd effect on mRNA level in the nine-day old barley and maize plants (Lysenko et al. 2015). The current experiment confirmed these results: mRNA levels of 13 chloroplast genes remained unchanged in both typical and atypical 16-day old Cd-treated barley plants (Fig. 10A). However, mRNA *psbD* decreased in Cd-treated plants (Fig. 10B). In the atypical plants, *psbD* level was always smaller than in the untreated plants, while the typical Cd-treated plants showed great diversity of variants. In the several biological experiments and/or cDNA synthesis, *psbD* in the typical Cd-treated plants was a) at the level of untreated plants, b) intermediate, c) at the level of atypical plants, and once it was d) below the level of atypical plants. Therefore, *psbD* decrease in the atypical plants was more established than in the typical Cd-treated plants. In nine-day old plants, mRNA *psbD* was not studied.

**Figure 10.**
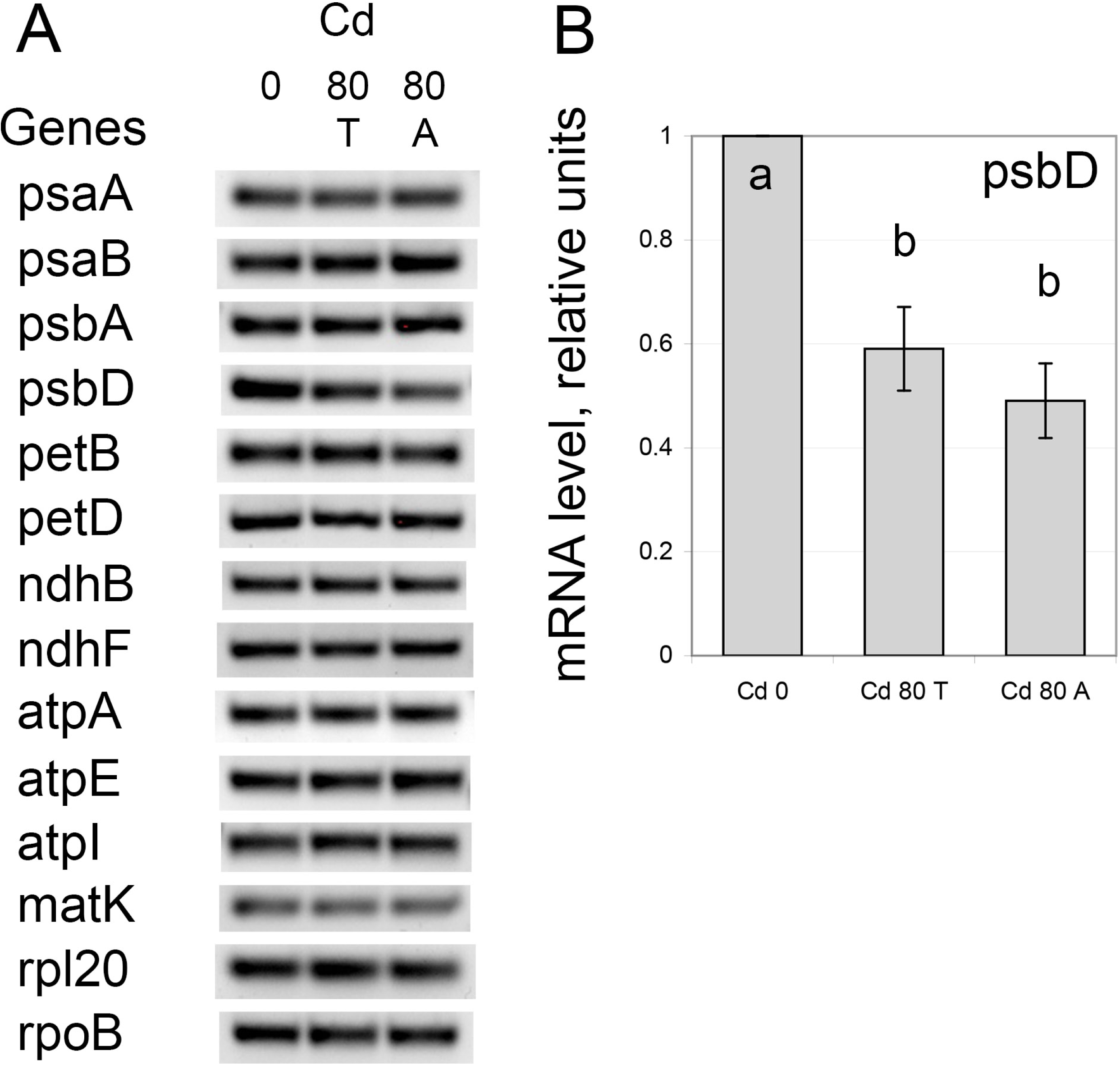
Levels of chloroplast mRNAs in the first leaves of 16-day old barley plants. A – PCR-products from mRNAs of 14 chloroplast genes. Typical results are shown, B – quantitative analysis of mRNA psbD. Means ± SE are shown. a-b – the differences are significant at p ≤ 0.05. Cd 0 – untreated plants, Cd 80 T – Cd-treated typical plants, Cd 80 A – Cd-treated atypical plants.

## 4. Discussion

Previously, we demonstrated a number of unusual Cd effects on the photosynthetic electron-transfer chain. They were as follows: A) in the dark-adapted state, the ratio Pm/Fv decreased; B) under illumination, the ratio Y(I)/X(II) decreased; C) qN decreased and D) qC increased (Lysenko et al. 2020). These effects obtained in nine-day old barley plants grown at 80 μM Cd; the effects were specific to Cd action (Lysenko et al. 2020). It would be more convincing to show that larger Cd toxicity enlarged these effects. We applied 250 μM Cd that is lethal in prolonged experiment for both barley (Lysenko et al. 2015) and maize (Klaus et al. 2013); however, we failed to show a larger effect at the higher Cd concentration. Probably, this caused by smaller Cd accumulation *in vivo*. Barley plants at 250 μM Cd, accumulated less Cd in leaves and, probably, chloroplasts than barley plants at 80 μM Cd (Lysenko et al. 2015). Maize plants also accumulated less Cd in chloroplasts comparing with barley counterparts (Lysenko et al. 2015); in addition, maize plants own C_4_-type of photosynthesis and Kranz anatomy. We used the alternative *in vivo* model to study the effects of higher Cd toxicity. We studied 16-day old barley plants grown at 80 μM Cd and compared the typical ones with the atypical more damaged plants from the same vessels. The analysis extended by the measurement of low temperature spectra of Chl fluorescence and stomatal conductance that we did not study before.

### 4.1. Balance of PSI : PSII

The dark-adapted plants show the maximal photochemical activity of RC PSII (Fv) and RC PSI (Pm); Fo is attributed to the antennae of both photosystems (Pfündel et al. 2013). Cd treatment decreased both Fv and Pm. At 80 μM Cd, Fv decreased slightly in both barley and maize; the decrease can be insignificant (Fig. 7C; Lysenko et al. 2020; Lysenko and Kusnetsov 2024) or significant (Figs. 1, 3B). Under higher Cd toxicity, Fv decreased more both at 250 μM Cd (Lysenko and Kusnetsov 2024) and in atypical plants at 80 μM Cd (Figs. 1, 3B, 7C) in both species; solely in Fig. 1, the decrease was insignificant because only three plants were measured. In the most cases, Pm decreased more than Fv that was demonstrated by decreases of the ratio Pm/Fv. At 80 μM Cd, Pm/Fv decreased significantly or insignificantly in the same experiments. In 9-day old maize plants, Pm/Fv decreased significantly (Fig. 1) or remained unchanged (Lysenko and Kusnetsov 2024); in 9-day old barley plants, Pm/Fv decreased significantly (Lysenko et al. 2020; Lysenko and Kusnetsov 2024); in 16-day old barley plants, Pm/Fv decreased significantly (Fig. 7E) or insignificantly (Fig. 3E). Under higher Cd toxicity, the decrease of Pm/Fv was always significant. In 9-day old plants, Pm/Fv similarly decreased in barley at 80 and 250 μM Cd (Lysenko and Kusnetsov 2024) and in the typical and atypical maize plants at 80 μM Cd (Fig. 1); in maize plants, Pm/Fv unchanged at 80 μM Cd and decreased at 250 μM Cd (Lysenko and Kusnetsov 2024). In 16-day old barley plants grown at 80 μM Cd, Pm/Fv in the atypical plants decreased more than in the typical ones (Figs. 3E, 7E).

All these data suggest that Cd decreased amount of the photochemically active PSI (Pm) more than amount of the photochemically active PSII (Fv); higher Cd toxicity enchanced this effect that was obvious in the atypical barley plants (Figs. 3E, 7E). The typical both untreated and Cd-treated barley plants showed weak or no correlation between Pm and Fv values; the distribution of Pm/Fv was rather cloud-like (Fig. 3F). In the atypical plants, Pm and Fv decreased proportional to each other demonstrating clear tendency (R^2^ = 0.877, Fig. 3F). Probably, the large loss of functionally active PSI drove the concomitant fall of functionally active PSII content.

Under illumination, Cd treatment unexpectedly increased the actual photochemical activity of PSII (X(II)) in barley plants. In IC, X(II) increased similarly at 80 and 250 μM Cd (9d, AL 128; Lysenko and Kusnetsov 2024) and at 80 μM Cd in both typical and atypical plants (16d, AL 168, Fig. 4A). In RLC, X(II) increased at 80 μM Cd in plants of both ages (Fig. 4B, Lysenko and Kusnetsov 2024); in more damaged plants; X(II) mostly remained at the control level (Fig. 4B, Lysenko and Kusnetsov 2024). Cd treatment decreased the actual photochemical activity of PSI (Y(I)) in 9-day old barley plants at 80 and 250 μM Cd (Lysenko and Kusnetsov 2024). In 16-day old barley plants, a different pattern was observed. In the typical plants, Y(I) increased in IC (Fig. 5A) and remained at the control level in RLC (Fig. 5B). In the atypical plants, Y(I) was unchanged or decreased (Fig. 5A–B).

At 80 μM Cd, the ratio Y(I)/X(II) decreased in barley plants of both ages (Fig. 6A–B), Lysenko and Kusnetsov 2024); thus, Cd treatment shifted the balance of actual activities of photosystems also in favor of PSII. In more damaged plants, we failed to accelerate this shift in 9-day old plants at 250 μM Cd (Lysenko and Kusnetsov 2024), while observed further decrease of Y(I)/X(II) in the atypical 16-day old plants (Fig. 6A–B). In maize, a pattern of X(II) and Y(I) changes was quite different (Lysenko and Kusnetsov 2024).

The low temperature spectra of Chl fluorescence are mainly applied for studies of antennae and process of state transition (Kalaji et al. 2014); however, we also studied Cd effect using this method. The normalized low temperature spectra demonstrated the opposite tendency: a shift in favor of PSI. In the typical Cd-treated plants, peak of PSI increased comparing with untreated plants (Fig. 7A). In the atypical plants, peak turned back to the level of untreated plants or even below it (Fig. 7A–B). The difference between Cd-treated typical and atypical plants shifted in more red area of the spectrum comparing with the difference between the typical untreated and Cd-treated plants (Fig. 7B).

Probably, Cd influenced fluorescence spectra by means of two independent factors. One factor increased PSI-fluorescence in the typical Cd-treated plants. In more damaged atypical plants, another factor was induced that decreased fluorescence. The similar pattern observed for X(II) at 80 μM Cd and in more damaged plants (see this subchapter). A similar pattern observed for non-photochemical quenching; we suggested two reversely acting factors (Lysenko and Kusnetsov 2024). A similar pattern observed for stomatal conductance and we demonstrated two different reasons (see chapters 3.7 and 4.4). We speculate that Cd treatment (typical) increased PSI fluorescence by means of increased antenna fluorescence; in more damaged plants, PSI fluorescence decreased due to the decrease of RC PSI.

### 4.2. Non-photochemical quenching and acceptor-side limitation of PSII

At 80 μM Cd, non-photochemical quenching decreased in the typical plants of both ages (Fig. 4C–D, Lysenko et al. 2020, Lysenko and Kusnetsov 2024). This increased a portion of closed RC PSII that excited by light and quenched neither photochemically, nor non-photochemically (Fig. 4E–F, Lysenko et al. 2020, Lysenko and Kusnetsov 2024). A portion of closed RC PSII can be calculated with the coefficients qC (Lysenko et al. 2020) or 1–qP. The coefficient qC shows balance between qC and qN (qC + qN + X(II) = 1, Lysenko et al. 2020), while 1–qP omit non-photochemical quenching from analysis. In IC, non-photochemical quenching is moderate therefore both qC and 1–qP show similar pictures (Suppl. Fig. S3). In RLC at rather high light intensity, non-photochemical quenching is large therefore qC and 1–qP demonstrated drastically different dynamics (Suppl. Fig. S3).

At 250 μM Cd, 9-day old barley plants demonstrated no further decrease of qN with the concomitant increase of qC (Lysenko and Kusnetsov 2024). While, the atypical 16-day old plants demonstrated larger decrease of qN and increase of qC comparing with the typical ones (Fig. 4C-F).

Both qC and Y(ND) changed actively. However, we revealed planes of the ratio qC/Y(ND) indicating that qC and Y(ND) changed proportional to each other and their ratio remained constant; these horizontal spans were similar in IC and RLC (Lysenko and Kusnetsov 2024). 16-day old barley plants demonstrated this feature much better. In the untreated plants, qC/Y(ND) remained at the level ∼0.2 in both IC and RLC (Fig. 6C). Cd-induced loss of non-photochemical quenching increased this level. In the typical Cd-treated plants, qC/Y(ND) remained at the level ∼0.4 in both IC and RLC (Fig. 6C). In the atypical Cd-treated plants, qC/Y(ND) further increased to the level 0.7-0.8 in IC; in RLC, the ratio qC/Y(ND) gradually decreased from 0.8 to 0.6 along with the increase of AL intensity (Fig. 6C). 16-day old barley plants kept the balance of limitations at the acceptor side of PSII and the donor side of PSI. Limitation at the acceptor side of PSII was smaller and did not limit the electron transfer between PSII and PSI. However, the untreated plants quenched non-photochemically large portion of qC. In the typical Cd-treated plants, this ability was reduced; in more damaged atypical plants, this ability was reduced more (Fig. 6C).

### 4.3. Metal contents

Which factor caused chlorosis in the atypical barley plants? First, we verified Cd content in the first leaves. The contents of Cd were similar in the typical 9-day old (Lysenko et al.2020) and 16-day old plants (Fig. 8A); also, Cd contents were similar in both typical and atypical 16-day old plants (Fig. 8A). Thus, increased Cd accumulation was not a reason for more damage. Unfortunately, we were limited in chlorotic leaves and did not determine Cd content in their chloroplasts.

K is the major metal in the first leaf of barley plants (Fig. 8D; Lysenko et al. 2020). A large set of soil-grown barley varieties demonstrated similar K accumulation in leaves: 1.0–1.6 mmol/g DW (Wu et al. 2015). In wheat leaves, K content was smaller by an order of magnitude (Ouzounidou et al. 1997).

Cd treatment decreased K content in the first leaves (Fig. 8D). In the atypical plants, K decreased even more than in the typical Cd-treated plants (Fig. 8D); this extra decrease was close to the significant level per g DW (p < 0.055) and significant per single leaf (Suppl. Table S3). This was the sole decrease of metal per gDW in the atypical plants; all the other metals per gDW increased in the atypical plants comparing with the typical Cd-treated plants whether significant or not (Fig. 8, Suppl. Table S3). The sum of Na, Mg, Ca, Fe, Zn, Mn, and Cu increased in the atypical plants (375 ± 28 μmol/g DW) significantly comparing with the untreated plants (302 ± 13 μmol/g DW); in the typical Cd-treated plants, the sum was even smaller but with larger dispersion (294 ± 37 μmol/g DW). Probably, the atypical plants suffered from severe K deprivation and increased general uptake of cations to compensate K shortage.

During the second week of barley growth, K content increased in the first leaves of untreated plants from 1.36 ± 0.12 mmol/g DW on the 9^th^ day (Lysenko et al. 2020) to 2.09 ± 0.04 mmol/g DW on the 16^th^ day (Suppl. Table S3); the difference is significant at p < 0.01 if different experiments can be compared. Whereas in Cd-treated plants, K contents slightly changed from the 9^th^ to 16^th^ day: 1.42 ± 0.10 (16d T) ≥ 1.29 ± 0.14 (9d T) ≥ 1.16 ± 0.08 (16d A) mmol/g DW. In a pilot experiment, barley plants with no Cd and K in the mineral media grown similar to a control until the 9th day; however, they were reduced and demonstrated signs of chlorosis on the 16^th^ day in contrast to a control (not shown). Probably, Cd treatment inhibited K uptake from the mineral media, while K content in seeds was sufficient for barley growth up to the 9^th^ day. This is why we did not observe K decrease in 9-day old plants (Lysenko et al. 2020). In the atypical plants, K uptake can be more severe inhibited and/or they have less K in their seeds. So, K deprivation can be the reason of Cd-induced chlorosis in the atypical plants.

Barley plants exchanged two major bivalent cations as the result of Cd treatment: Mg content increased, Ca contents decreased, while their sum remained unchanged (Fig. 8C, Suppl. Table S3). Cd treatment decreased the content of Cu. The contents of Na, Fe, Mn, and Zn did not change significantly (Suppl. Table S3).

### 4.4. Stomatal conductance

K is a key driver of stomatal movement (Shimazaki et al. 2007). Therefore, we studied stomatal conductance in barley plants. Once again, the stomatal conductance in the typical Cd-treated plants appeared to be higher than in both untreated plants and atypical Cd-treated plants (Fig. 9A). We were surprised because Cd inhibition of stomatal conductance is well documented. Cd treatment decreased stomatal conductance (Shi and Cai 2008), stomatal density (Baryla et al. 2001), stomata openness (Barcelo et al. 1988), and conductivity of a single stoma (Zhu et al. 2005). To unravel this contradiction, we studied diurnal dynamics of stomatal conductance in the typical plants; we were unable studying a diurnal dynamic in the atypical plants due to their small number.

The untreated and Cd-treated barley plants demonstrated different patterns of stomatal conductance over a daytime. The untreated plants kept stomatal conductance at the adaxial (upper) side higher than at the abaxial (lower) side of leaf blade (Fig. 9B). The untreated plants fulfilled complicated diurnal dynamics of stomatal conductance including both opening and closing. We have no idea why the untreated barley plants demonstrated such unusual pattern of g_sw_. However, unusual diurnal dynamics of g_sw_ reported in some plant species grown in a greenhouse under natural light conditions (Suwannarut et al. 2023).

The typical Cd-treated plants did not differentiate stomatal conductance at adaxial and abaxial sides of leaves; the difference was small and insignificant (Fig. 9B). The typical Cd-treated plants performed simple diurnal dynamics with a plateau or slow increase until 14.00 and a subsequent gradual decrease. As a whole, stomatal conductance was equal in the untreated and typical Cd-treated plants during all the daytime (Fig. 9C); the mean values comprise the data from 8.00 to 21.00 at both sides of leaves. We suppose that K shortage hindered stomatal movement. Therefore, the typical Cd-treated plants avoided extreme closing and opening to keep sufficient stomatal conductance during the daytime. At 10.00, g_sw_ was low in the untreated plants because they are flexible in regulation of stomatal movement; g_sw_ was higher in the typical Cd-treated plants because they are less flexible in the regulation (Fig. 9A). In the atypical Cd-treated plants, further K shortage decreased stomatal conductance comparing with the typical Cd-treated plant. We suppose that it was the overall decrease of g_sw_ while we were able measuring it at 10.00 solely.

### 4.5. Chloroplast mRNA levels

Previously, we demonstrated that Cd treatment did not influence mRNA levels of 10 chloroplast genes; it was demonstrated in 9-day old barley and maize plants grown at 80 and 250 μM Cd (Lysenko et al. 2015). The current study supported earlier conclusion: mRNA levels of 13 chloroplast genes were changed neither in typical nor in atypical Cd treated plants at 80 μM Cd (Fig. 10A). However, mRNA *psbD* decreased in both Cd-treated variants (Fig. 10 B). In previous study, *psbD* was not analyzed (Lysenko et al. 2015). Therefore, we cannot conclude that it was age-specific decrease.

In chloroplasts, *psbD* is highly transcribed (*e.g.*, Zubo et al. 2008) and highly regulated at the transcriptional level (*e.g.*, Thum et al. 2001). Heat stress drastically decreased the maximal activity of RC PSII (Fv) in barley plants (Lysenko et al. 2023); this Fv decrease was larger than under Cd-induced stress (Fig. 3B, 7C). Under heat stress, we observed concerted activation of transcription of *psbA, psbD, psaA,* and *psaB* in barley chloroplasts (Zubo et al. 2008); these genes code for D1 and D2 subunits of RC PSII and A1 and A2 subunits of RC PSI correspondingly. Probably, transcription of these genes has been activated to sustain activities of both photosystems under heat stress. In Cd-treated barley plants, the sole decrease of *psbD* transcript is dissimilar to the concerted reaction observed under the heat stress. The chloroplast gene expression machinery is located in stroma. In stroma, Cd content is low; the most part of Cd translocated to thylakoids (Lysenko et al. 2019). The sole decrease of *psbD* transcript looks like a result of indirect Cd action. Perhaps, *psbD* decrease resulted from general stress conditions with no especial damage to RC PSII.

## Abbreviations

AL: actinic light
Chl: chlorophyll
IC: induction curve
OEC: oxygen-evolving complex
PSI / PSII: photosystem I / II
RLC: rapid light curve
RC: reaction center
SP: saturation pulse
g_sw_: stomatal conductance

## PAM terms

Fv/Fm – maximal quantum yield of PSII

ΔF – peak induced with SP reflecting portion of photochemically active PSII

ΔC – Chl fluorescence induced with AL reflecting portion of closed PSII

qP, Փ_PSII_, X(II) – photochemical coefficients of PSII

qN – coefficient of non-photochemical quenching of excited Chl state

qC – coefficient of acceptor-side limitation of PSII (closed PSII)

Y(I) – photochemical coefficient of PSI

Y(ND) – coefficient of donor-side limitation of PSI

Y(NA) – coefficient of acceptor-side limitation of PSI

Fo / Fo’ – minimal Chl fluorescence in dark / light, induced with measuring light

Fm / Fm’ – maximal Chl fluorescence in dark / light

Fv / Fv’ – variable Chl fluorescence in dark / light

Fs – stationary Chl fluorescence in light (synonym F)

Po – minimal light absorption by P_700_

Pm / Pm’ – maximal P_700_ change (P_700_°/ P_700_^+^) in dark / light

P – stationary P_700_ light absorption level in light

## Author contribution

**Eugene A. Lysenko**: Conceptualization, Methodology, Project administration, Investigation, Visualization, Formal Analysis, Funding Acquisition, Writing; **Natalia A. Savvina:** Investigation, Formal Analysis; **Alexander V. Kartashov**: Investigation; **Galina V. Kochetova**: Methodology, Investigation. All authors read and approved the manuscript. EAL is the author responsible for contact and ensures communication.

## Acknowledgement

We are grateful to Drs L.A. Koppel for access to the fluorometer and M.A. Breygina for providing liquid nitrogen.

