## Supplemental Tables and Figures for "Atypical chlorotic plants as a tool for studying more severe Cd effect on photosystem I, non-photochemical quenching, K content, and stomatal conductance"

**Source:** bioRxiv

Table S1. List of primers and other conditions of RT-PCR.

| Gene | Primers (5' → 3') | Product size, bp | Annealing temperature, ºC | Number of cycles |
|---|---|---|---|---|
| psaA | S1: GGACCCAGTGCTCAGGTAGTT<br>AS1: TGGGAGCGGCTTTGTGATA | 226 | 60 | 20-22 |
| psaB | S1: ACCCCACTACTCGTCGTATTTG<br>AS1: CACTGATAAACCCCAGAATAAGC | 325 | 60 | 20-24 |
| psbA | S1: GATGGTTCGGTGTTTTGATGAT<br>AS1: CCACTCACGACCCATATAACAAG | 302 | 60 | 14-19 |
| psbD | S1: ACCCAACTCAAGCTGAAGAAAC<br>AS1: CTGCCATCCACGCACGAATA | 285 | 60 | 18-21 |
| petB | S: GGTATCTCTGGAATATGAGT<br>AS: CATCATACTTGCTGACCAT | 293* | 58 | 25-26 |
| petD | S: AAGGCGGATTATGGGAGT<br>AS: CAATACCTAACCAAAGAGCTAC | 446* | 63 | 20-23 |
| ndhB | S2: GGCAAGCTCTTCTATTCTGGT<br>AS: TGAGTAATAGCAAGGAGATTCC | 387* | 63 | 25-27 |
| ndhF | S2: CCTCTTCTCCCACTTCCAGTTA<br>AS2: CAAATATCCTTCATCGTGAGACAT | 333 | 60 | 25-28 |
| atpA | S2: AGTATGACCGCTTTACCAATAGT<br>AS2: CCTTGCCAATTGATTCTGACTT | 303 | 63 | 21-23 |
| atpE | S: CGAATTATTTGGGATTGTGAAGTG<br>AS: TCGTATCCTAGCTCGTCTGAGAG | 345 | 60 | 24-26 |
| atpI | S: TCATAGCTGTTCGGAATCCAC<br>AS: CTCAATCCTTTTTTGCTAAGACC | 313 | 60 | 24-26 |
| matK | S2: CGGGGCATCCTATTAGTAAAC<br>AS2: CCGAACCCAATCGTTGCATA | 222 | 63 | 25-26 |
| rpl20 | S: TCGTCGTTTGTGGATCACT<br>AS: ATCGTGTAAAGATTATTTGGA | 163 | 58 | 26-30 |
| rpoB | S2: GTCCTGGTATTTACTACCGCT<br>AS: GTCGCATAAGGACTCCGAA | 350 | 63 | 27-33 |

* - from spliced mRNA.

Tables S2. The portion of atypical plants with signs of chlorosis among 16-day-old barley plants grown at $CdS0_4$ 80 μM.

| Experiment | All plants | Atypical plants with signs of chlorosis * | |
|---|---|---|---|
| 1 | 204 | 7 | 3.43% |
| 2 | 165 | 10 | 6.06% |
| 3 | 216 | 7 | 3.24% |
| 4 | 217 | 5 | 2.30% |
| 5 | 53 | 4 | 7.55% |
| 6 | 220 | 5 | 2.27% |
| 7 | 220 | 3 | 1.36% |
| 8 | 220 | 5 | 2.27% |
| 9 | 220 | 4 | 1.82% |
| 10 | 191 | 2 | 1.05% |
| 11 | 220 | 4 | 1.82% |
| 12 | 191 | 5 | 2.62% |
| 13 | 110 | 2 | 1.82% |
| Sum | 2447 | 63 | |
| Mean ± SE | | | 2.9 ± 0.5% |

* – plants with yellowish shade at the first leaf were chosen and tested with Dual-PAM-100; those demonstrating the ratio Fv/Pm ≥ 3.8 were selected as atypical.

The threshold level 3.8 selected empirically. In the early experiments, all the typical Cd-treated plants showed the ratio Fv/Pm < 3.6. Out of the whole 76 tested typical Cd-treated plants, three only showed the ratio Fv/Pm = 3.9-4.12.

We were limited with the number of atypical plants for the analyses of low temperature Chl fluorescence and RNA content. In these experiments, the plants with slight signs of chlorosis and Fv/Pm < 3.8 were also used. These 10 plants are not included in the Table S1.

Table S3. Metal contents in the first leaves of 16-day-old barley plants.

| Metal | µmol / g DW | | | nmol / leaf | | |
|---|---|---|---|---|---|---|
| | Cd 0 | Cd 80 T | Cd 80 A | Cd 0 | Cd 80 T | Cd 80 A |
| Cd | $0.00 \pm 0.00^{b}$ | $1.82 \pm 0.17^{a}$ | $2.26 \pm 0.37^{a}$ | $0.0 \pm 0.0^{b}$ | $21.8 \pm 1.6^{a}$ | $20.1 \pm 3.4^{a}$ |
| K | $2\,090 \pm 40^{a}$ | $1\,421 \pm 95^{b}$ * | $1\,159 \pm 75^{b}$ * | $27\,121 \pm 1080^{a}$ | $16\,814 \pm 1532^{b}$ | $10\,026 \pm 939^{c}$ |
| Na | $175 \pm 15^{a}$ | $167 \pm 28^{a}$ | $218 \pm 17^{a}$ | $2\,194 \pm 198^{a}$ | $1\,976 \pm 277^{a}$ | $1\,930 \pm 181^{a}$ |
| Mg | $48 \pm 1^{b}$ | $78 \pm 8^{a}$ | $95 \pm 8^{a}$ | $604 \pm 21^{b}$ | $926 \pm 69^{a}$ | $842 \pm 88^{a}$ |
| Ca | $76 \pm 3^{a}$ | $46 \pm 2^{c}$ | $58 \pm 4^{b}$ | $957 \pm 44^{a}$ | $555 \pm 28^{b}$ | $517 \pm 46^{b}$ |
| Mg+Ca | $125 \pm 2^{b}$ | $124 \pm 9^{ab}$ * | $153 \pm 11^{a}$ * | $1561 \pm 47^{a}$ | $1481 \pm 62^{a}$ | $1359 \pm 127^{a}$ |
| Fe | $1.63 \pm 0.06^{a}$ | $1.69 \pm 0.13^{a}$ | $1.70 \pm 0.08^{a}$ | $20.4 \pm 1.0^{a}$ | $20.5 \pm 1.9^{a}$ | $15.0 \pm 0.9^{b}$ |
| Mn | $0.762 \pm 0.093^{a}$ | $0.625 \pm 0.028^{a}$ | $0.718 \pm 0.037^{a}$ | $9.50 \pm 1.06^{a}$ | $7.51 \pm 0.17^{a}$ | $6.34 \pm 0.41^{b}$ |
| Zn | $0.280 \pm 0.054^{a}$ | $0.242 \pm 0.035^{a}$ | $0.331 \pm 0.044^{a}$ | $3.54 \pm 0.75^{a}$ | $2.89 \pm 0.37^{a}$ | $2.96 \pm 0.46^{a}$ |
| Cu | $0.110 \pm 0.009^{a}$ | $0.062 \pm 0.002^{c}$ | $0.073 \pm 0.004^{b}$ | $1.383 \pm 0.140^{a}$ | $0.749 \pm 0.025^{b}$ | $0.648 \pm 0.053^{b}$ |

Cd 0 – untreated plants, Cd 80 T – Cd-treated typical plants, Cd 80 A - Cd-treated atypical plants.

Cd treatment – 80 µM.

Mean ± SE.

a-c – difference is significant at $p < 0.05$.

Color fonts visualize significant differences to ease perception.

* – difference between the metal contents in Cd80 T and Cd80 A leaves is significant at:

K – at $p < 0.055$. Mg+Ca – at $p < 0.065$.

Figure S1. The first leaves of 16-day old barley plants.
An experiment with the most severe damaged atypical leaves.

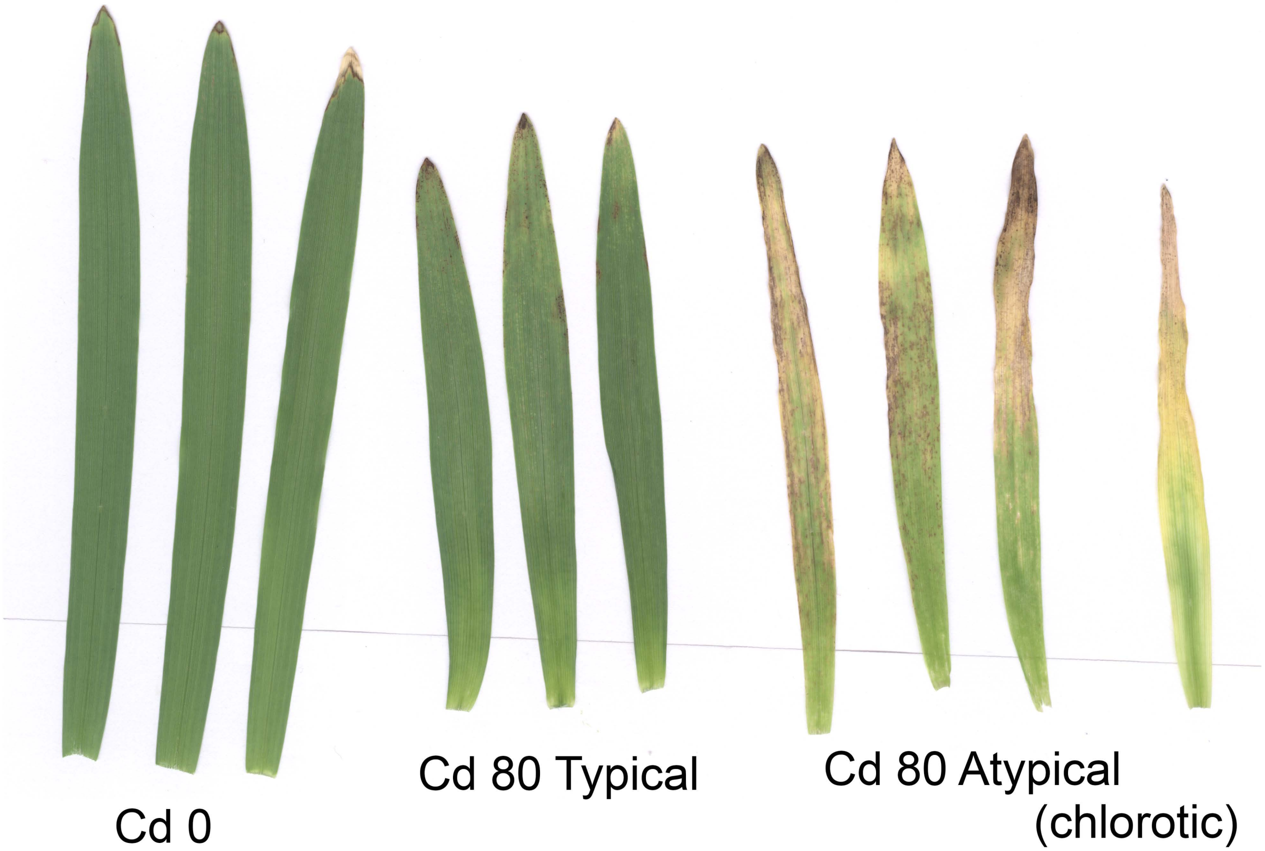


General remarks:
In 16-day old barley plants, necrosis did not influence the ratio Pm/Fv, while chlorosis (yellow color or yellowish shade) did.

A completely yellow first leaf (with no necrosis) was obtained once out of ~ 2 500 Cd-treated plants studied (in other experiment, not shown).

Figure S2. The photochemical part ΔF of modulated Chl fluorescence and ΔF-based photochemical coefficients .

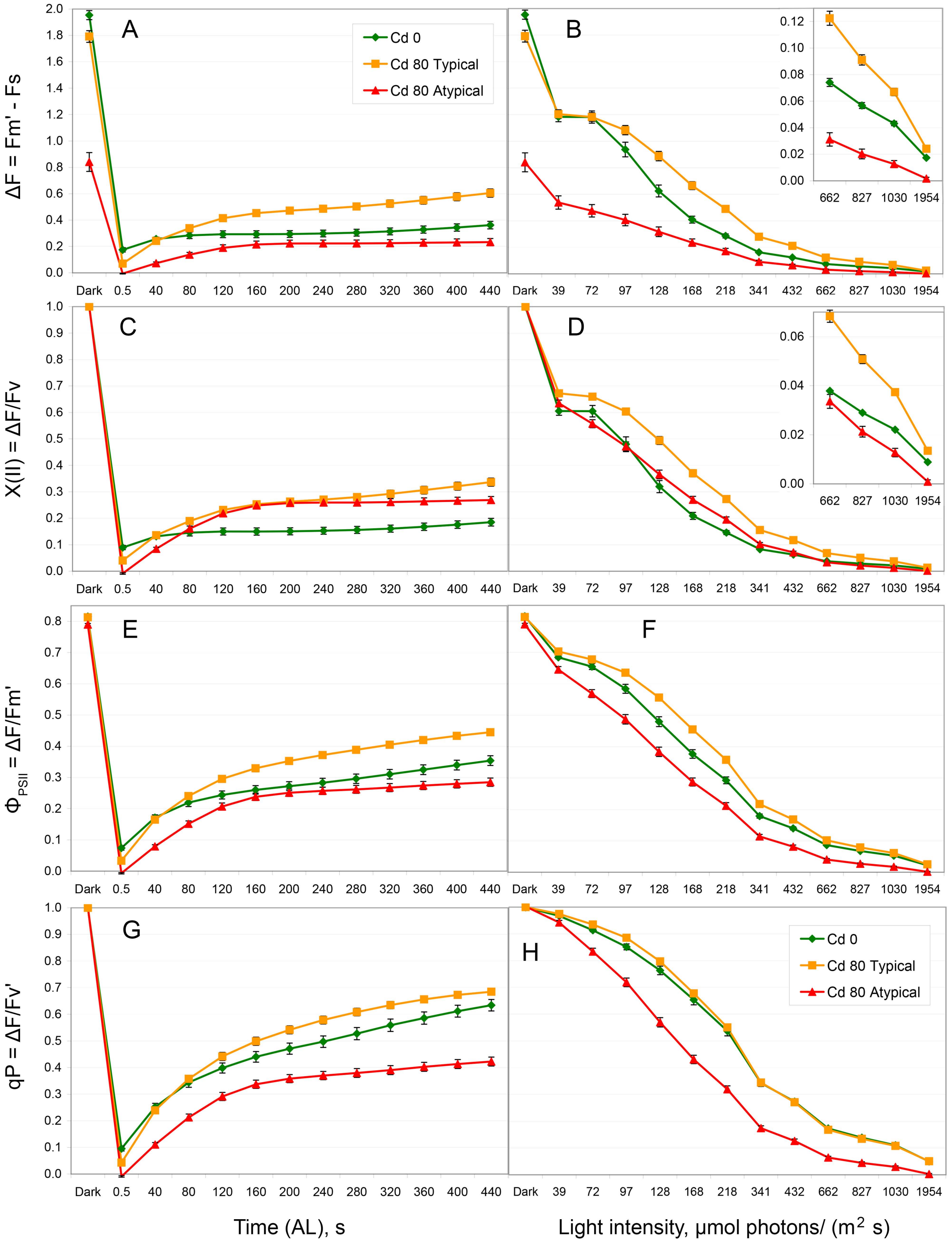


ΔF = Fm' – Fs, a peak induced with SP under an AL illumination.

A-B – ΔF; C-D – X(II); E-F – ΦPSII; G-H – qP. A, C, E, G – IC; B, D, F, H – RLC.
Green diamonds – untreated plants (Cd 0), orange squares – Cd-treated typical plants (Cd 80 Typical), red triangles – Cd-treated atypical plants (Cd 80 Atypical).
Means ± SE are shown. The insets show the corresponding small values with the higher resolution.

Figure S3. Limitation of PSII at the acceptor side (closed PSII): primary values (ΔC) and the coefficients qC and 1–qP.

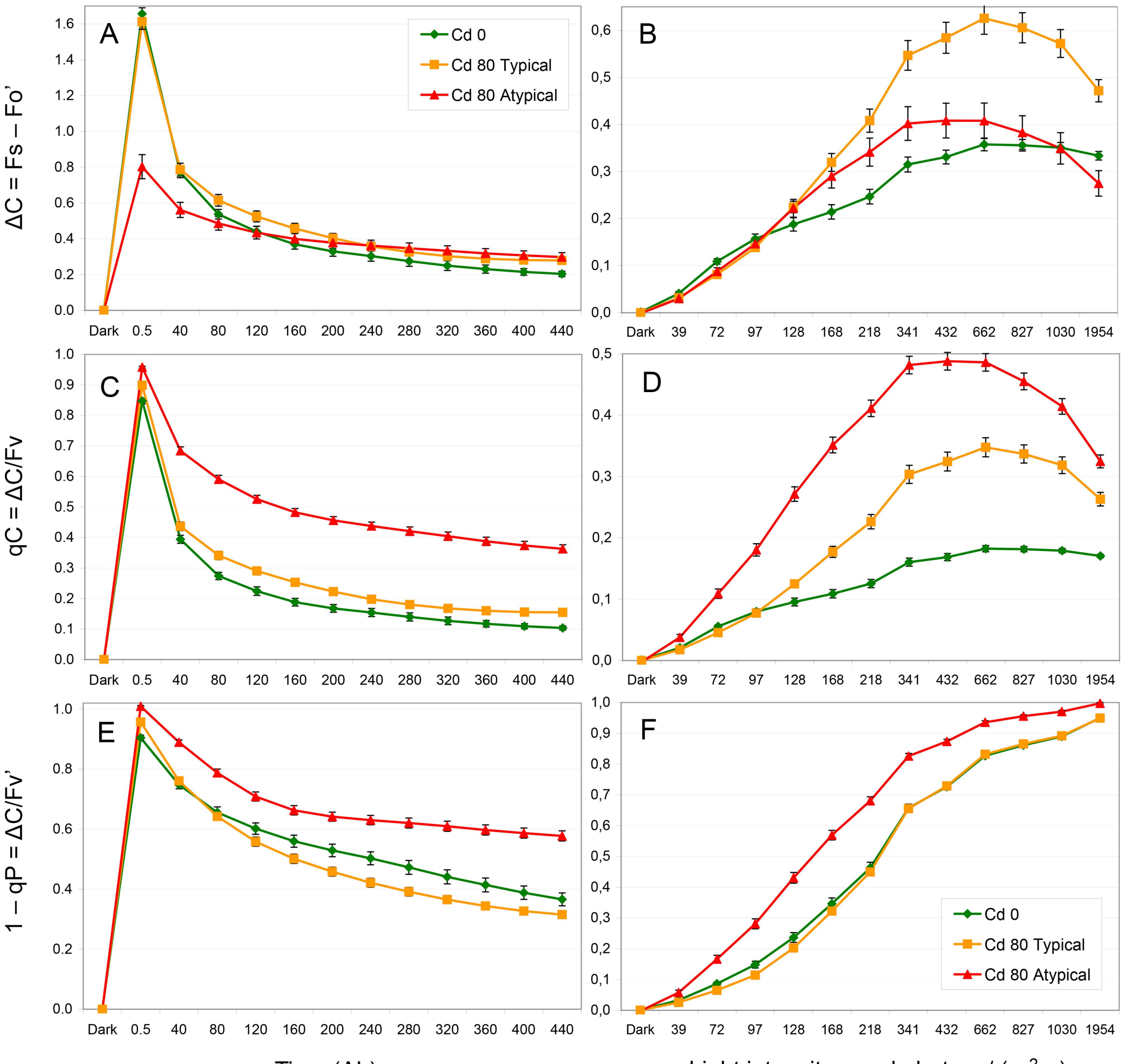


ΔC = Fs –Fo', a fluorescence induced with an AL illumination.

A-B – ΔC; C-D – qC; E-F – 1–qP. A, C, E – IC; B, D, F – RLC.
Green diamonds – untreated plants (Cd 0), orange squares – Cd-treated typical plants (Cd 80 Typical), red triangles –- Cd-treated atypical plants (Cd 80 Atypical).
Means ± SE are shown. The insets show the corresponding small values with the higher resolution.

Figure S4. Maximal activities of PSII (Fv) and PSI (Pm) determined in seven experiments prior to the measurements of low temperature Chl fluorescence or RNA content.

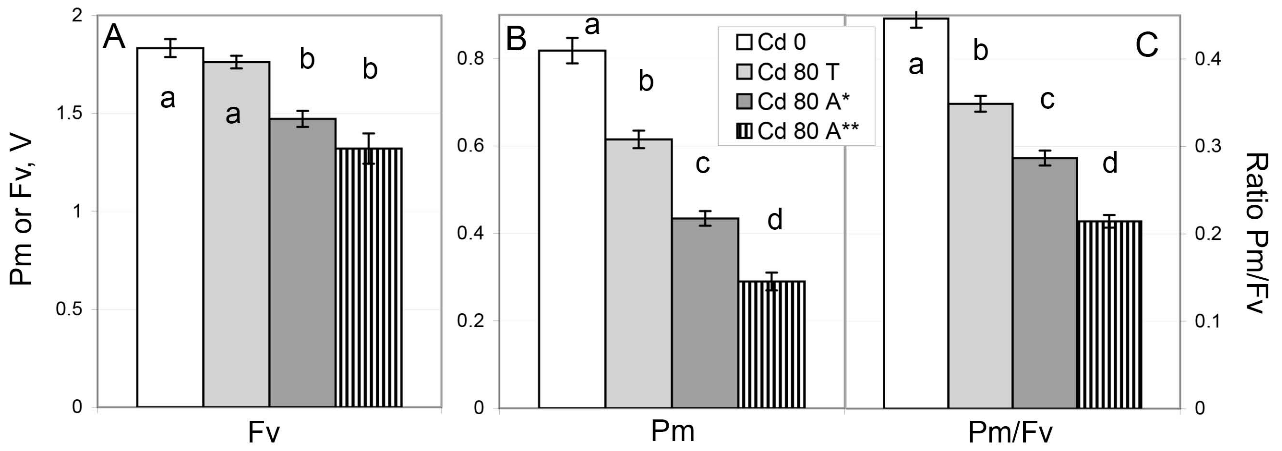


A – Fv; B – Pm; C – Pm/Fv.
White bars – untreated plants (Cd 0).
Light grey bars – Cd-treated typical plants (Cd 80 T).
Dark grey bars – Cd-treated plants primary selected by the phenotype (Cd 80 A*).
Vertically hatched bars – Cd-treated plants selected by the phenotype and further demonstrated $Fv/Pm \geq 3.8$ (Cd 80 A**).
Means ± SE are shown.
a-d – the differences are significant at $p \leq 0.05$.

$Fv/Pm = 3.8$ corresponds to $Pm/Fv = 0.263$
